# The first Miocene fossils of Lacerta cf. *trilineata* (Squamata, Lacertidae) with a comparative study of the main cranial osteological differences in green lizards and their relatives

**DOI:** 10.1101/612572

**Authors:** Andrej Čerňanský, Elena V. Syromyatnikova

## Abstract

We here describe the first fossil remains of a green lizardof the *Lacerta* group from the late Miocene (MN 13) of the Solnechnodolsk locality in southern European Russia. This region of Europe is crucial for our understanding of the paleobiogeography and evolution of these middle-sized lizards. Although this clade has a broad geographical distribution across the continent today, its presence in the fossil record has only rarely been reported. In contrast to that, the material described here is abundant, consists of a premaxilla, maxillae, frontals, parietals, jugals, quadrate, pterygoids, dentaries and vertebrae. The comparison of these elements to all extant green lizard species shows that these fossils are indistinguishable from *Lacerta trilineata*. Thus, they form the first potential evidence of the occurrence of this species in the Miocene. This may be also used as a potential calibration point for further studies. Together with other lizard fossils, Solnechnodolsk shows an interesting combination of survivors and the dawn of modern species. This locality provides important evidence for the transition of an archaic Miocene world to the modern diversity of lizards in Europe. In addition, this article represents a contribution to the knowledge of the comparative osteological anatomy of the selected cranial elements in lacertid. This study gives special emphasis to the green lizards, but new data are also presented for related taxa, e.g., *Timon lepidus, Podarcis muralis* or *Zootoca vivipara*. Although the green lizards include several cryptic species for which determination based on isolated osteological material would be expected to be difficult, our comparisons show several important morphological differences.

## Introduction

Although the fossil record of squamates is well documented in western and central Europe, many aspects of eastern Europe remain unknown. However, this area can be crucial for our understanding of the biogeography and evolution of this currently dominant group of non-avian reptiles [1–3]. We here describe a significant find of fossil remains that can be allocated to the lizard family Lacertidae from Upper Miocene beds near ther town of Solnechnodolsk (Fig. 1; the Ponto-Caspian steppe region) in southern European Russia - MN 13 of the European Neogene Mammal biochronological system [4]. The lizard clade Lacertidae includes over 300 small-to medium-sized living species [5], which are broadly distributed in Eurasia and Africa. Lacertids form the dominant reptile group in Europe, where this group is also suggested to have originated [6]. This is supported by the fossil record [7–9]. The Lacertidae forms a monophyletic group [10] consisting of two lineages: Lacertinae and Gallotiinae [6]. Descendants of the basalmost divergence in Lacertidae, between Gallotiinae and Lacertinae, are also documented from Europe [11–12]. The members of the clade Lacertidae exhibit an extensive range of body sizes. The medium-sized lacertids, belonging to Lacertinae, are represented by the green lizards of the genus *Lacerta*. Molecular and morphological phylogenetic analyses show that *Lacerta* is the sister taxon to *Timon* [13–17, 6].

**Fig 1.**
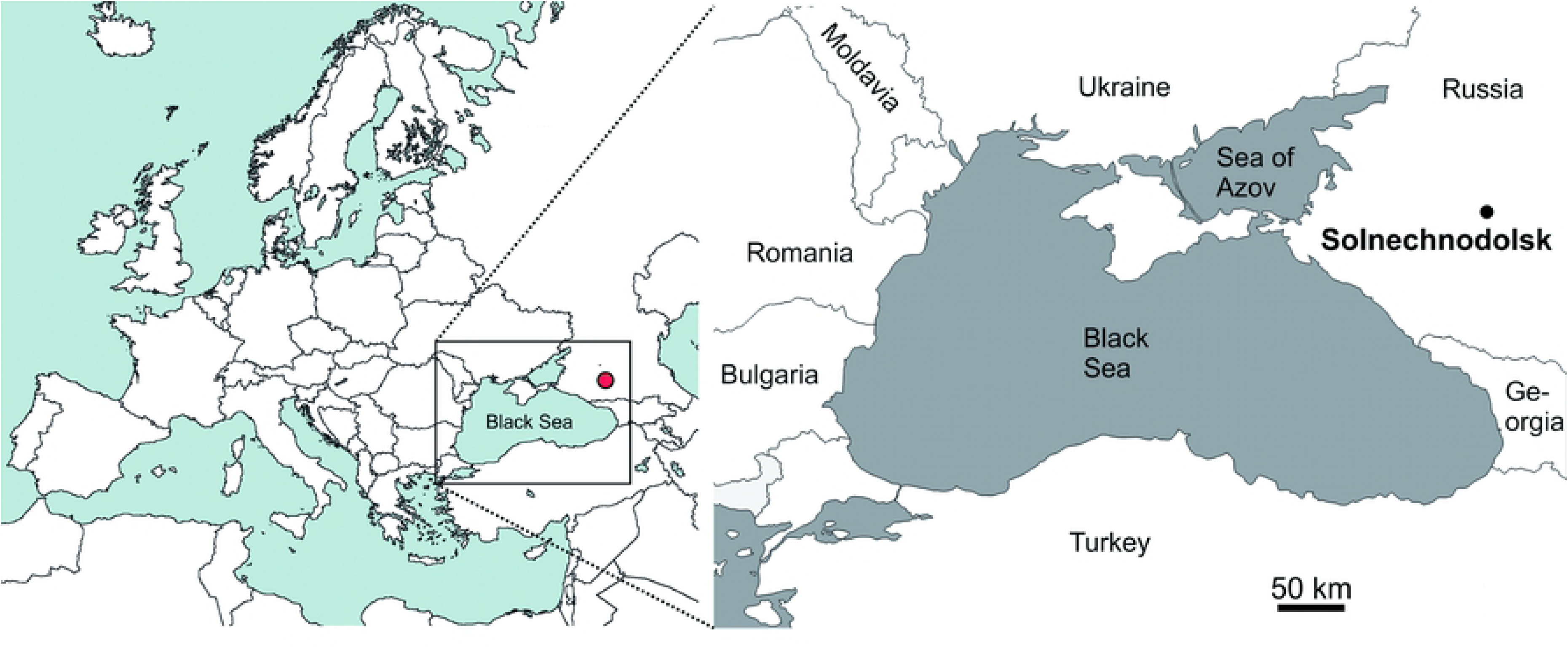
The location of the locality Solnechnodolsk.

The oldest finds of green lizards are documented from the lower Miocene (MN 4) locality Dolnice near Cheb in the Czech Republic [18]. The oldest known remains potentionally allocated to *Lacerta* cf.*viridis* have been described on the basis of several isolated jaw elements from the late Miocene (MN 11) localities of Kohfidish in Austria [19]. and Polgárdi in Hungary [20]. Nowodays, green lizards inhabit a large area extending from the Iberian Atlantic coast to Central Asia [21]. However, this group includes several morphologically cryptic species [22] for which determination based on isolated osteological material has been extremely difficult [18]. Even though there is useful published data concerning the osteology of these animals, it is not sufficiently detailed to allow reliable interpretation of fossil remains to species level. Peters [23] already recognized five species: *L. agilis, L. schreiberi* (Schreiber’s green lizard or the Iberian Emerald lizard), *L. strigata* (the Caucasus green lizard), *L. trilineata* (Balkan green lizard or three-lined green lizard) and *L. viridis*. Based on morphological studies [24], *L. trilineata* was recently split into three species, *L. trilineata, L. media* (Levant green lizard) and *L. pamphylica* (Turkisch pamphylic green lizard). Within *Lacerta viridis* clade, two main lineages were recognized - Western European lineage of *L. bilineata* and Eastern European *L. viridis* [25]. The taxonomy, evolution and biogeography of these taxa has been highly discussed [21, 16, 25–27]. However, the fossil record of green lizards from the crucial time period - Miocene, when the speciation within the group very likely occurred [16], is almost absent. This leads to a gap of our knowledge of the evolution of this group and to the absence of calibration points, resulting in wide confidence intervals for divergence time estimates [16, 27–28]. An essential requirement for enabling progress in interpreting the fossil record of *Lacerta* is a comparative study of extant taxa for the purpose of discovering characters useful in species level identification.

The Solnechnodolsk locality (45°18’N, 41°33’E) is situated in the Northern Caucasus, 40 km NW of the city of Stavropol. In paleogeographical terms, it is located at the southern shore of the Pontian marine basin. The vertebrate fossils occurred as disassociated bones coming from fluviatile and lacustrine beds incised in Middle Sarmatian (Bessarabian) limestones. Only preliminary accounts of the fauna have been so far published [29–31]. The data obtained from small mammals correlates the Solnechnodolsk fauna with the late Turolian. The locality yielded one of the most abundant and diverse vertebrate faunas of the late Miocene in Russia. Among them, only remains of *Pelobates*, amphisbaenians and anguimorphs have recently been described [3, 32]. In addition, the first Neogene mabuyid skinks are also documented from this site [33].

The aims of this paper are as follows: (1) to describe the fossil lacertid material from the late Miocene of Solnechnodolsk in detail; (2) to compare it with other lacertids, with the main focus on extant green lizards; (3) to do a comparative anatomy of selected cranial elements useful for diagnostic purposes in extant lizards. This will serve as a reference for other comparative anatomical studies of any other lacertids, extant, or extinct (such a study is crucial for thedetermination of the abundant record of the fossil lacertid cranial elements).

## Material & Methods

### Specimens examined

The lizard specimens described here are housed in the Geological Institute of the Russian Academy of Sciences, Moscow, Russia, prefixed under individual GIN numbers. Since the site discovery in 2009, the material was sampled in 2009, 2010, 2014, and 2017 by expeditions from of the Russian Academy of Sciences. Fossils were extracted from sediments by screen washing with the mesh size 0.5, 0.7, and 1 mm. The excavated vertebrate material includes fishes, amphibians, reptiles, birds, small and large mammals.

### Specimens used for comparisons

The following specimens of extant lizard species were used for comparison: *Lacerta viridis* (NHMV 40137, DE 51, 131, 132), *L. schreiberi* (NHMV 10809), *L. bilineata* (NHMV 18599), *L. pamphylica* (ZSM 1047/2005 - male; ZSM 939/2005 - female, NHMV 35861 - juvenile), *L. strigata* (NHMV 39765), *L. trilineata* (NHMV 27665; UF 65017), *L. media* (NHMV 34808), *L. agilis* (NHMV 39028), *Timon lepidus* (NHMV 10921-1), *Podarcis muralis* (NHMV 39359-1), *Zootoca vivipara* (NHMV 32438-1), *Takydromus sexlineatus* (pers. coll. of A. Č.), *Meroles ctenodactylus* (NHMV 31376); *Psammodromus algirus* (NHMV36038-2) and *Gallotia stehlini* (NHMV 11031-1).

#### Institutional abbreviations

DE: Department of Ecology, Comenius University in Bratislava, Slovakia;
GIN: Geological Institute of the Russian Academy of Sciences, Moscow, Russia;
NHMV: Museum of Natural History, Vienna;
UF: University of Florida, Florida Museum of Natural History (USA);
UMJGP: Universalmuseum Joanneum, Graz, Austria;
ZSM: The Bavarian State Collection of Zoology, Munich, Germany.

### X-ray microtomography, three-dimensional visualization and photography

The specimens of extant lacertids were scanned by the micro-computed tomography (CT) facility at the Slovak Academy of Sciences in Banská Bystrica, using a a Phoenix mikro-CTv|tome|x L240. The CT data sets were analyzed using VG Studio Max 3.2 and Avizo 8.1 on a high-end computer workstation at the Department of Ecology (Comenius University in Bratislava). The fossil specimens and the maxilla of *Lacerta viridis* (DE 51) were photographed under a Leica M125 binocular microscope with axially mounted DFC 500 camera [LAS software (Leica Application Suite) version 4.1.0 (build 1264)].

### Anatomical terminology, character reconstruction and measurements

The standard anatomical orientation system is used throughout this paper. The image processing program ImageJ [34] was used for measurements. We used Mesquite (Version 2.75; [35]) to optimize characters using parsimony. The Mesquite metatree is based on 26 Marzahn et al. [26], Kornilios et al. [27], Pyron et al. [15] and Čerňanský & Smith [9] for lacertids and outgroup relationships.

### Systematic paleontology

Squamata [36]

Lacertoidea [37]

Lacertidae [36]

*Lacerta* [38]

***Lacerta* cf. *trilineata*** [39]

Figs. 2–7

**Fig 2.**
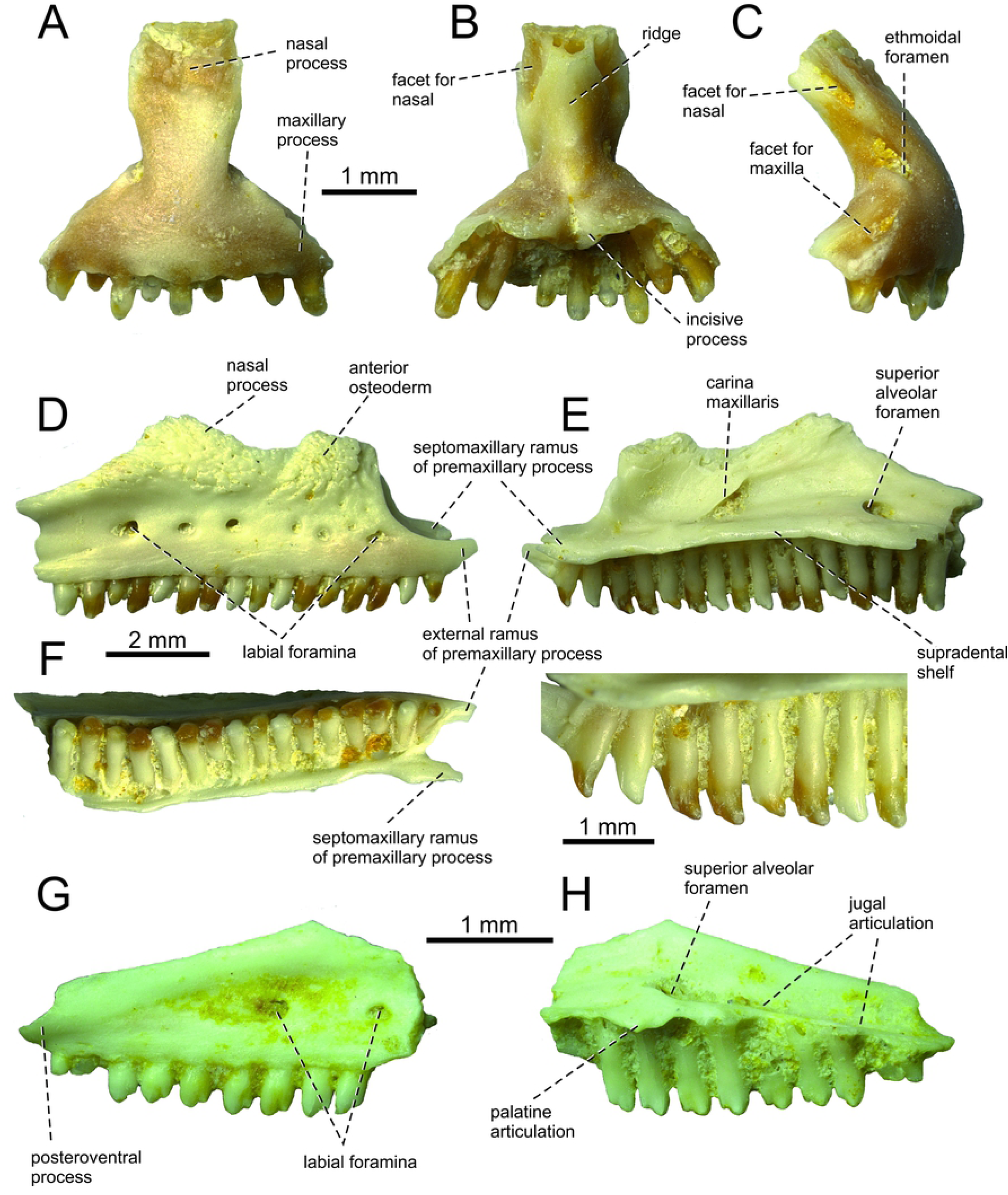
*Lacerta* cf. *trilineata* from the late Miocene of the locality Solnechnodolsk. Premaxilla GIN 1145/270 in external (A), internal (B), and lateral aspects (C). Maxilla GIN 1145/271 in lateral (D), medial (E) with a detail of teeth, and ventral aspects (F). Maxilla GIN 1145/272 in lateral (G), and medial aspects (H).

Material––One premaxilla GIN 1145/270, six right maxillae 1145/271-276, five right frontals 1145/277-281, one left frontal 1145/282, four parietals 1145/283-285, 304, one left prefrontal 1145/286, two right jugals 1145/287-288, one left jugal 1145/289, two right postfrontals 1145/290-291, one right quadrate 1145/292, one right pterygoid 1145/293, two left dentaries 1145/294-295, four right dentaries 1145/296-299.

Locality and horizon––Solnechnodolsk (Stavropol Region, southwestern Russia); upper Miocene, MN 13.

## Results

### Osteological Description

#### Premaxilla

The premaxilla is a triradiate element (Fig. 2A-C). The nasal process is broad, gradually widening posterodorsally. Here, the internal surface bears the facet for the nasal on each side. Between these facets, a distinct ridge is developed. The external surface of the nasal process bears ornamentation. Unfortunately, apex of this process is broken off. The portion below the nasal process is roughly triangular in shape. The maxillary processes are well developed, but short. On their lateral edges, they bear a wedge shaped facet for the premaxillary process of the maxilla on each side. The dental portion bears nine tooth positions (eight teeth are still attached). The subdental shelf forms a short, slightly bilobed incisive process in its mid-region. The vomerine processes is well developed.

#### Maxilla

The description is based on the specimen GIN 1145/271 (Fig. 2D-F), which represents a right maxilla. It bears 16 tooth positions - all teeth are still attached. However, the posterior portion of the maxilla is broken off, so the tooth number of the complete tooth row is certainly slightly higher. The supradental shelf (sensu Rage & Augé [40]) is thin, well expanded medially and slightly dorsally convex. The anterior extremity of the maxilla is divided into a short external ramus, the premaxillary process, and a broader, more medially oriented and slightly more dorsally developed septomaxillary (internal) ramus. The orientation of the septomaxillary ramus is anterior rather than being dorsally curved as it is in *Timon lepidus* or *Zootoca vivipar*a [41]. An oval premaxillary fenestra is present between these rami. Immediatelly dorsally to this fenestra, a small foramen is present –the anterior opening for superior alveolar canal. The nasal process is broad, but its dorsal portion is damaged. This process bears the carina maxillaris (sensu Müller [42]; this structure passes between the vestibulum and the cavum nasi, see [43]), which starts to rise posterodorsally from the level of the 6th tooth position (counted from anterior). The superior alveolar foramen is located at the level between of the 13-14th. tooth positions (counted from anterior). If we use this position as a reference point in the specimen GIN 1145/272, the complete posterior portion of a maxilla (Fig. 2G-H), we can estimate 18-19 tooth positions in the complete tooth row.

The lateral face of the maxilla is pierced by 6 larger labial foramina arranged in a single line and several smaller, scattered in the anterior region. The posteriormost one of the larger series is located at the level between the 13th and 14th tooth position (counted from anterior) in 1145/271, or at the level of 7th tooth position (counted from posterior) in 1145/272. The nasal process bears attached osteodermal shields. At least two well separated osteoderms fused to lateral side of the ventral portion of the nasal process of maxilla can be identified here. They are seaparated by a sulcus. This sulcus has a posteroventral course and virtually points to the 4th labial foramen (counted from anterior). The sculpture of these osteoderms consists of densely spaced grooves, pits and ridges. The anteriorly located osteoderm is large. The length of its ventral margin forms 1/3 of the entire length of the sculptured region.The posteroventral process of maxilla does not gradually narrow to its end, but it is stepped. The notch between the dorsal portion and ventral portion is shallow, weakly developed. The posteroventral process rises anteriorly, gradually continues to the nasal process. In their contact, only small dorsal curvature is present (it indicates the posterior end of the nasal process).

#### Prefrontal

The left prefrontal is preserved (Fig. 3A-C). It is roughtly triradiate element. It has a long posterodorsal process, the ventral surface of which forms the anterodorsal margin of the orbit. In medial aspect, a facet for frontal can be observed. The ventral process, broader than the posterodorsal process, forms the anterior surface of the orbit. The dorsal surface possesses an ornamented osteodermal shield (the prefrontal shield). The ornamentation consists of pits, grooves and small ridges. The internal surface is deeply excavated for the nasal capsule. The dorsal region forms the contact with the nasal and maxilla.

**Fig 3.**
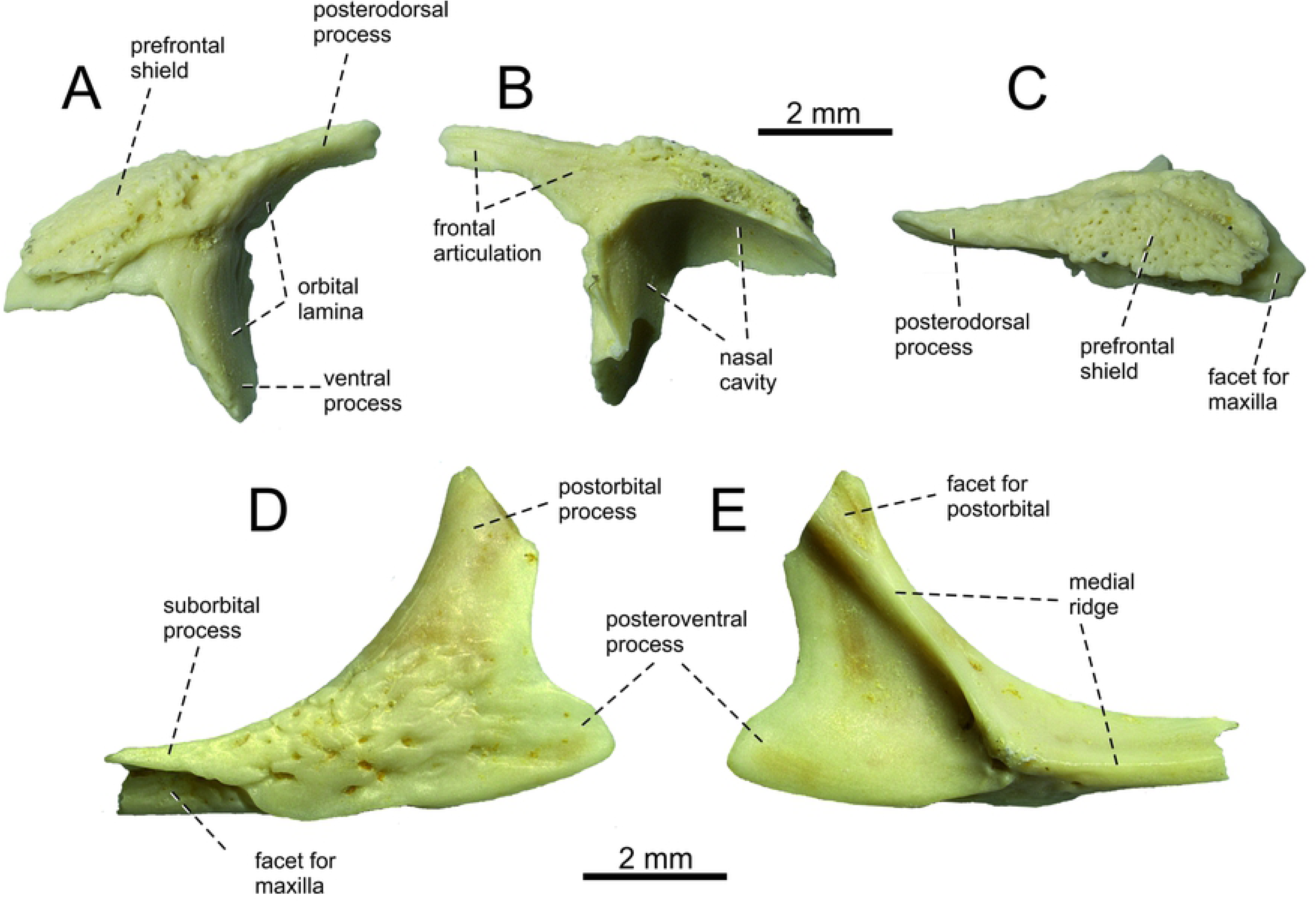
*Lacerta* cf. *trilineata* from the late Miocene of the locality Solnechnodolsk. Left prefrontal GIN 1145/286 in lateral (A), medial (B), and dorsal aspects (C). Left jugal GIN 1145/289 in lateral (D), and medial aspects (E).

#### Jugal

A right jugal is preserved but incomplete (Fig. 3D-E). It is a triradiate element. Only the ventral portion of the broad postorbital process is preserved - its dorsal termination is broken off. The posteroventral process is well defined. It is robust in lateral (or medial) aspect, roughtly trianglular in shape. The angle between the central line of the posteroventral and postorbital process of jugal is around 78°. The osteodermal shield is attached to the dorsolateral surface, mostly present in the suborbital process. Although the anterior portion of the suborbital process is not preserved, the posterior section of the facet for maxilla is visible in lateral aspect. In internal side, the medial ridge is present of the Type 1 in [44].

#### Frontal

There are six frontal bones preserved in the material. The description is mainly based on almost complete specimen GIN 1145/277 (Fig. 4A-C), which represents the right frontal. It is anteroposteriorly elogate element, roughtly rectangular in shape, with the posterior section being laterally expanded. The whole dorsal surface is covered by ornamented osteodermal shields, which are fused to the bone. Only exception is present in the anterior region, where a broad semi-elliptical smooth surface is located. It forms the articular facet for nasal. Laterally to it, a wedge-shaped smooth surface is located, representing the facet for the posterodorsal portion of the nasal process of the maxilla. The ornamentation of the osteodermal shields is indentical to that of the prefrontal and maxilla. It should be noted that in the specimen 1145/277, the prefrontal shield is less defined, although it is possible to distinguish it from the frontal shield. The anteroposterior maximum length of this frontal is 10 mm. The ornamentation is not so strongly developed as it is in larger specimens and thus we suggest that it represents a younger (immature) individual. In the specimen 1145/278 (Fig. 4D), the oval prefrontal shield can be easily recognized. The anterior end of the frontal is preserved only in GIN 1145/279. Here, the pointed anterolateral process is prominent, triangular in shape (Fig. 4E). Alhough only the base of the anteromedial process is preserved, it can be estimated (based on the size of the preseved portion) that it did not reach the level of the anterolateral process anteriorly. The sulcus interfacialis is well developed in several specimens, being slightly convex anteriorly. It forms a border between the frontal and frontoparietal shield. The lateral margin is slightly constricted anteriorly to the sulcus. The anterior region of the frontal in front of the sulcus interfacialis is much longer than the posterior region. It forms almost 2/3 of the entire anteroposterior length of the element. The contact with the parietal is interdigitated. The posterior region is especially well developed in 1145/282, where even the frontal portion of the parietal shield is preserved (see Fig. 4F). It occupies a small area on the posterolateral corner of the frontal.

**Fig 4.**
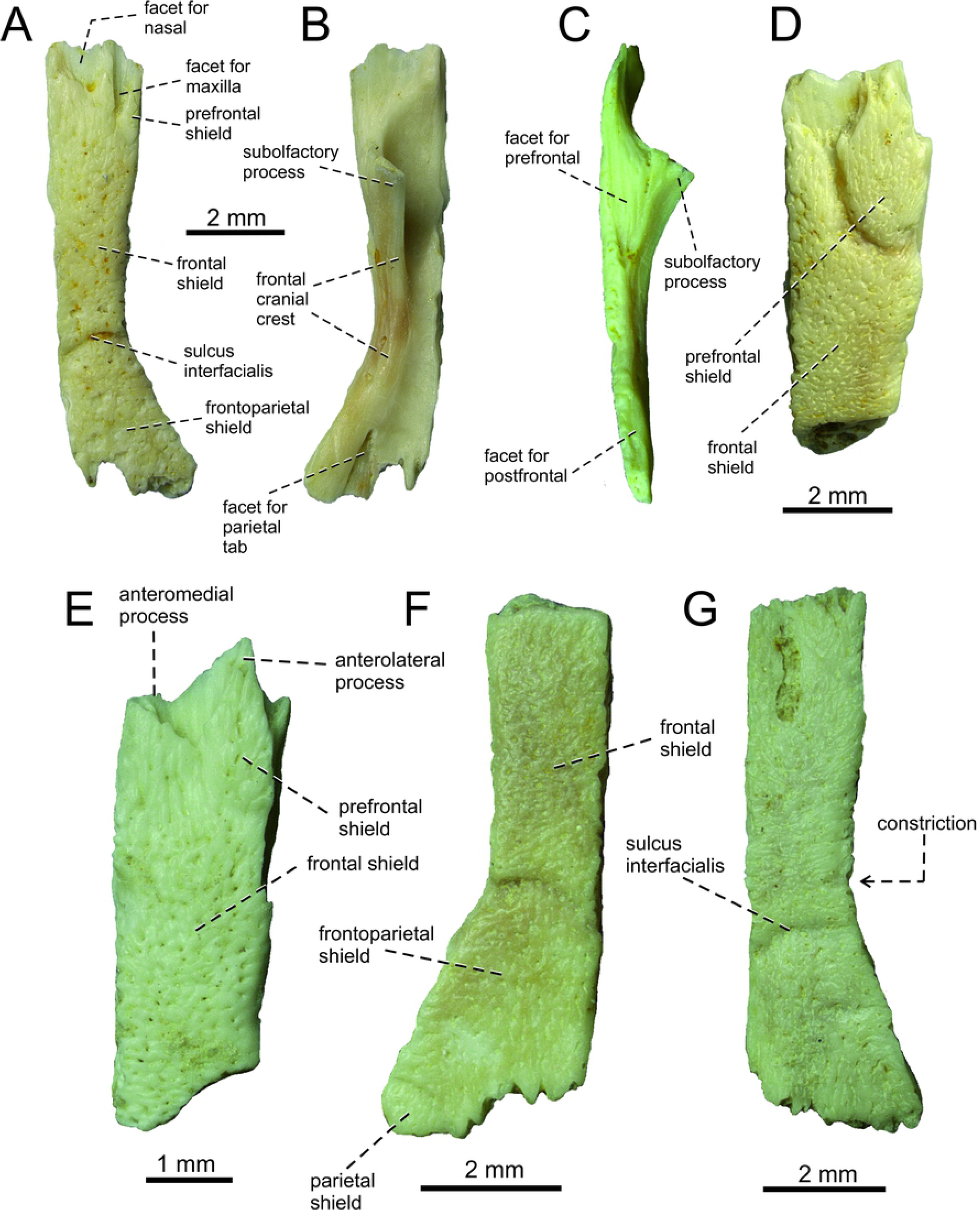
*Lacerta* cf. *trilineata* from the late Miocene of the locality Solnechnodolsk. Right frontals GIN 1145/277 (A-C), 1145/278 (D), 1145/279 (E), 1145/280 (G) and one lef frontal 1145/282 (F) in dorsal (A, D, E, F, G), ventral (B), and lateral (C) aspects.

On the lateral margin, a wedge-shaped broad facet for the prefrontal and a thin one for the postfrontal are not in contact, leaving the frontal exposed on the orbital margin. On the ventral side the frontal cranial crest forms the lateral orbital margin. It posseses two foramina in the posterior half of its length. The cranial crest expands anteriorly into a deep and well-developed subolfactory process. On the posterior margin is a well-developed triangular facet for the parietal tab.

#### Parietal

All parietals are incompletely preserved (Fig. 5). The description is mainly based on the specimens GIN 1145/283 (Fig. 5A-B) and 1145/284 (Fig. 5C-D). The parietal table is covered by several osteodermal shields. The centrally located interparietal shield is pierced by a parietal foramen in its anterior half. The interparietal shield is large, anteroposteriorly prolonged and pentagonal in shape. It is longer than a posteriorly located trapezoidal occipital shield. The occipital shield is, howerver, distinctly broad. It occupies a large area of the posterior portion of the partietal table. Laterally located parietal shields appear to be large, however they are not completely preserved. Only the right one shows the lateral margin, which appears to be slightly concave medially. The frontoparietal shields are only partly preserved on the left side in 1145/283. However, this region is better preserved in 1145/285, showing that the interparietal shield was entirely restricted to the parietal bone (Fig. 5E). The ornamentation of shields in parietals is identical to that of prefrontal and frontal. Only the base of the left supratemporal process is preserved in the specimen 1145/284. It is markedly broad.

**Fig 5.**
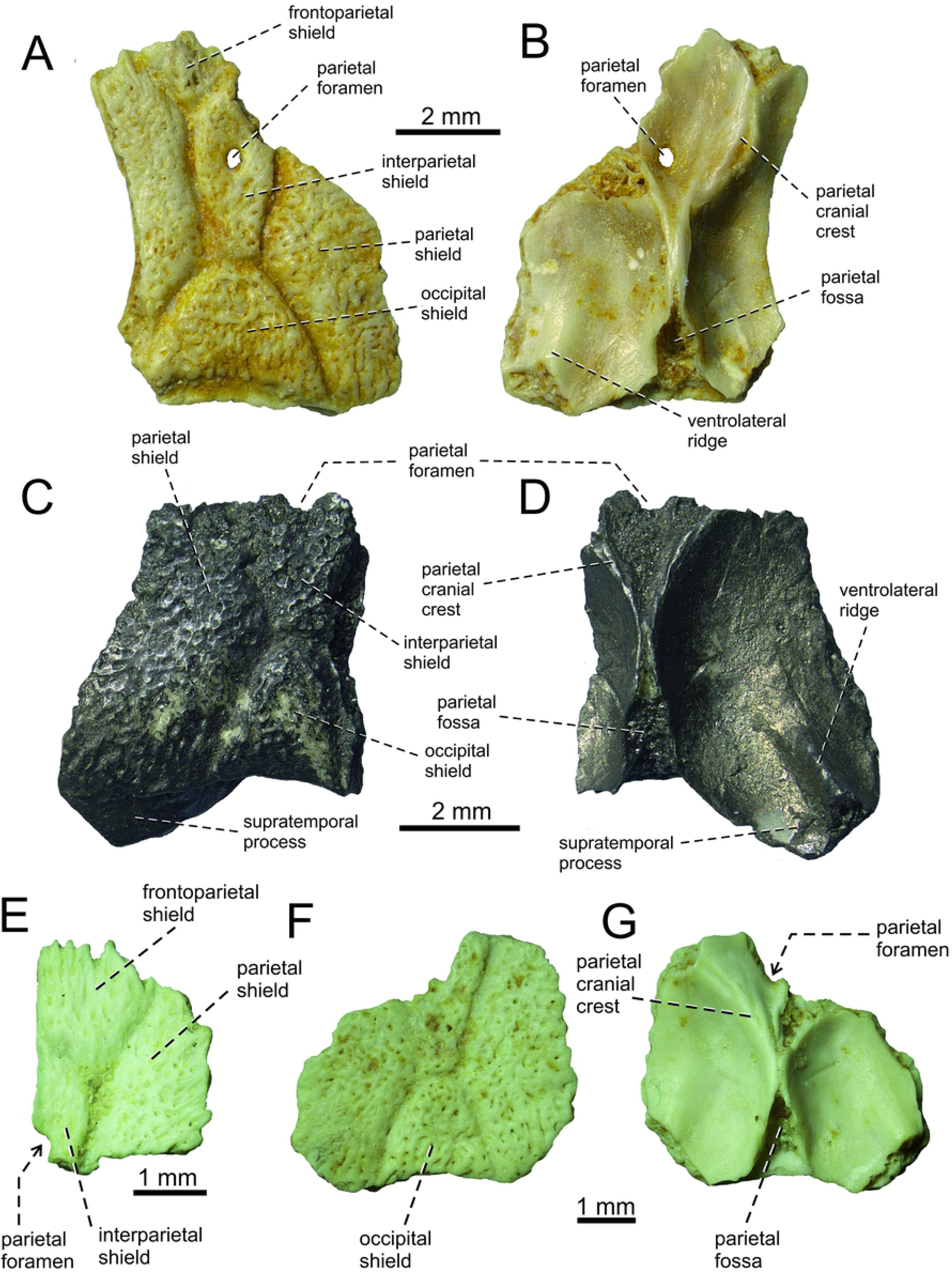
*Lacerta* cf. *trilineata* from the late Miocene of the locality Solnechnodolsk. Parietals GIN 1145/283 (A, B), 1145/284 (C, D), 1145/284, 1145/285 (E), and 1145/304 (F, G) in dorsal (A, C, E, F) and ventral (B, D, G) aspects.

On the ventral surface, parietal cranial crests are well-developed. They originate from the anterolateral corners of the parietal and converge posteromedially. The best preserved region bearing an anterior section of the parietal cranial crest can be seen in 1145/283 (Fig. 5B). Its course is not straight but it suddenly turns more medially approximately at the level of the anterior margin of the parietal foramen. Further posteriorly, the cranial crests merge posteriorly to the parietal foramen. They diminish posteriorly but continue to the end of the bone. A parietal fossa is located between them on the posterior midline (Fig. 5G). The ventrolateral ridge is developed in the lateral half of the supratemporal process here. It gradually disappears anteromedially. As mentioned, only the base of the supratemporal process is preserved and so the morphology of its distal portion is unknown.

#### Postfrontal

It is sub-rectangular and anteroposteriorly elongated, but its posterior section is damaged in both specimens (Fig. 6A-B). It bears anteroventral and anterodorsal expansions formed by the narrow, pointed postorbital process and a broad and blunt frontal process. This latter process bears a readily observed frontal articulation. The external surface of the postfrontal is mostly covered by an osteodermal shield. The internal surface is smooth, having a wedge shaped facet for an articulation with the postorbital.

**Fig 6.**
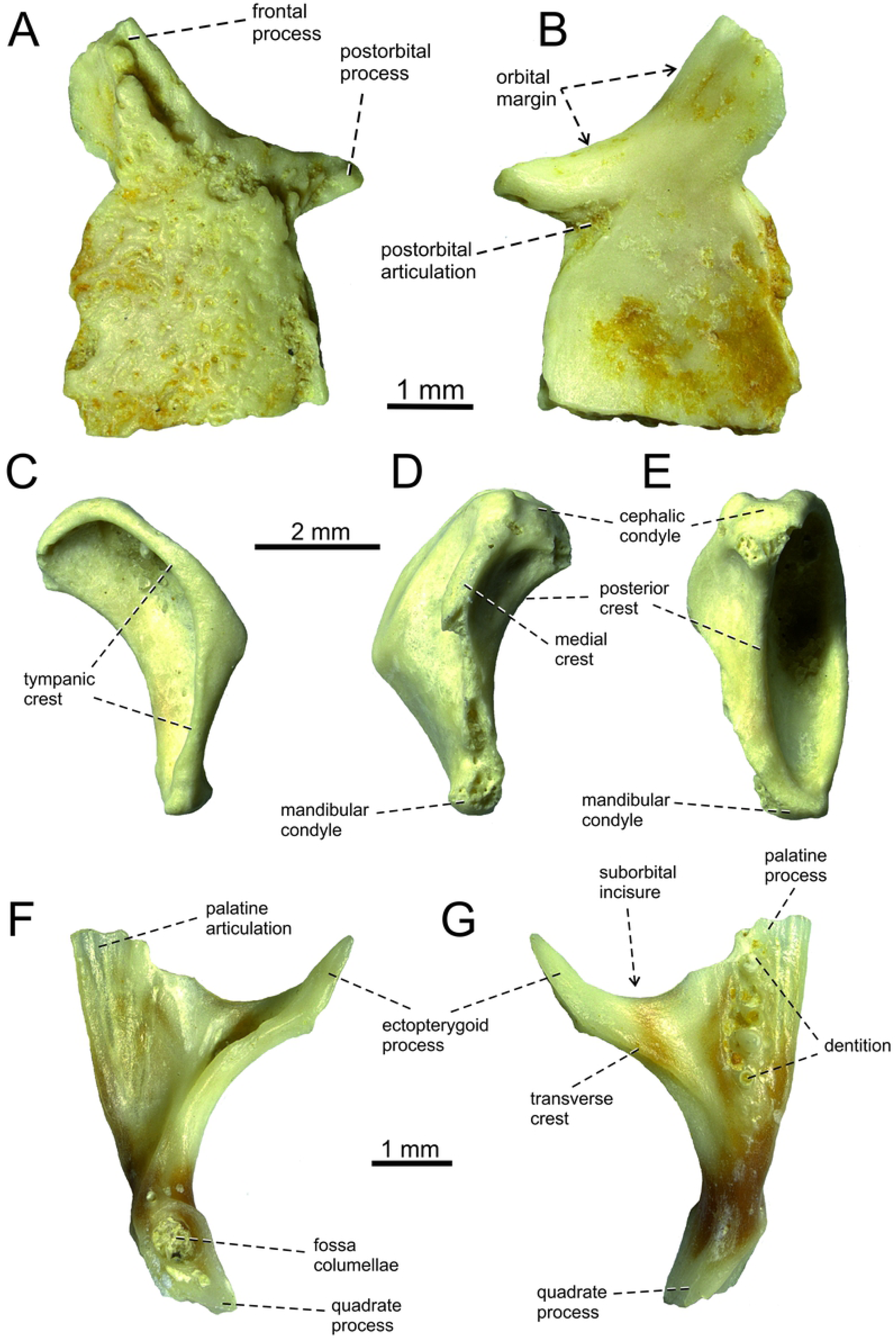
*Lacerta* cf. *trilineata* from the late Miocene of the locality Solnechnodolsk. Postfrontal GIN 1145/290 in external (A), and internal (B) aspects. Right quadrate GIN 1145/292 in lateral (C), medial (D), and posterior (E) aspects. Right pterygoid in dorsal (F), and ventral (G) aspects.

#### Quadrate

The quadrate is dorsoventrally elongated, narrow rather than robust (Fig. 6C-E). In lateral view, the quadrate is anteroposteriorly narrow, with an anteriorly expanded anterior margin. The anterior section is bordered by a laterally expanded sharp ridge: the tympanic crest. This crest is continuous from the cephalic to the mandibular condyle. The crest is angled approximately in the mid-region. The dorsal surface of the cephalic condyle protrudes slightly posteriorly. In lateral aspects, the ventral half of the quadrate gradually markedly narrows ventrally. Thus the ventral portion is much narrower than the dorsal portion. This ventral region ends with the saddle-shaped mandibular condyle. It is slightly smaller than the cephalic condyle. The medial surface posseses a distinct medial crest, however, only its dorsal section is completely preserved, whereas the ventral section is damaged.

#### Pterygoid

The incomplete right pterygoid is preserved (Fig. 6F-G). It is a tri-radiate, nearly letter’’y’’-shaped element. The ectopterygoid process is well developed, thin with a pointed anterior end. On the ventral surface, there is a transverse crest that runs along the base of the ectopterygoid process. The palatine process is broader, but its anterior end is broken off. The palatine process bears teeth on its ventral side, located in a single line. On the dorsal side, the palatine articulation is present. Between the palatine process and the ectopterygoid process, there is a broad area - the suborbital incisure. Only a small portion of the quadrate process is preserved. The fossa columellae (= epipterygoid fossa) is large, well defined by its rounded margin.

#### Dentary

Several dentaries are preserved, but all only incompletely (Fig. 7). The specimens GIN 1145/294 and 1145/296 represent only the anterior portions of the dentaries (see Fig. 7A-B). The most completely preserved specimen is 1145/295 (Fig. 7C-E). In this specimen, 21 tooth positions (15 teeth are still attached) are preserved. The subdental shelf (sensu Rage & Augé [40]) is rounded, dorsally concave. The symphyseal region is dorsally elevated and the symphysis is small, rectangular in shape. The subdental shelf gradually narrows posteriorly, mostly caused by the presence of the splenial here (the articulation with the splenial is still visible on the ventromedial side of the shelf, reaching the level of the 9th. tooth position if counted from anterior). Meckel’s groove is largely widely open, especially in the posterior direction. The alveolar foramen is located at the level between the 18th-19th tooth positions (counted from anterior in the specimen 1145/295, or 7th tooth position counted from posterior in 1145/297; see Fig. 7H).

**Fig 7.**
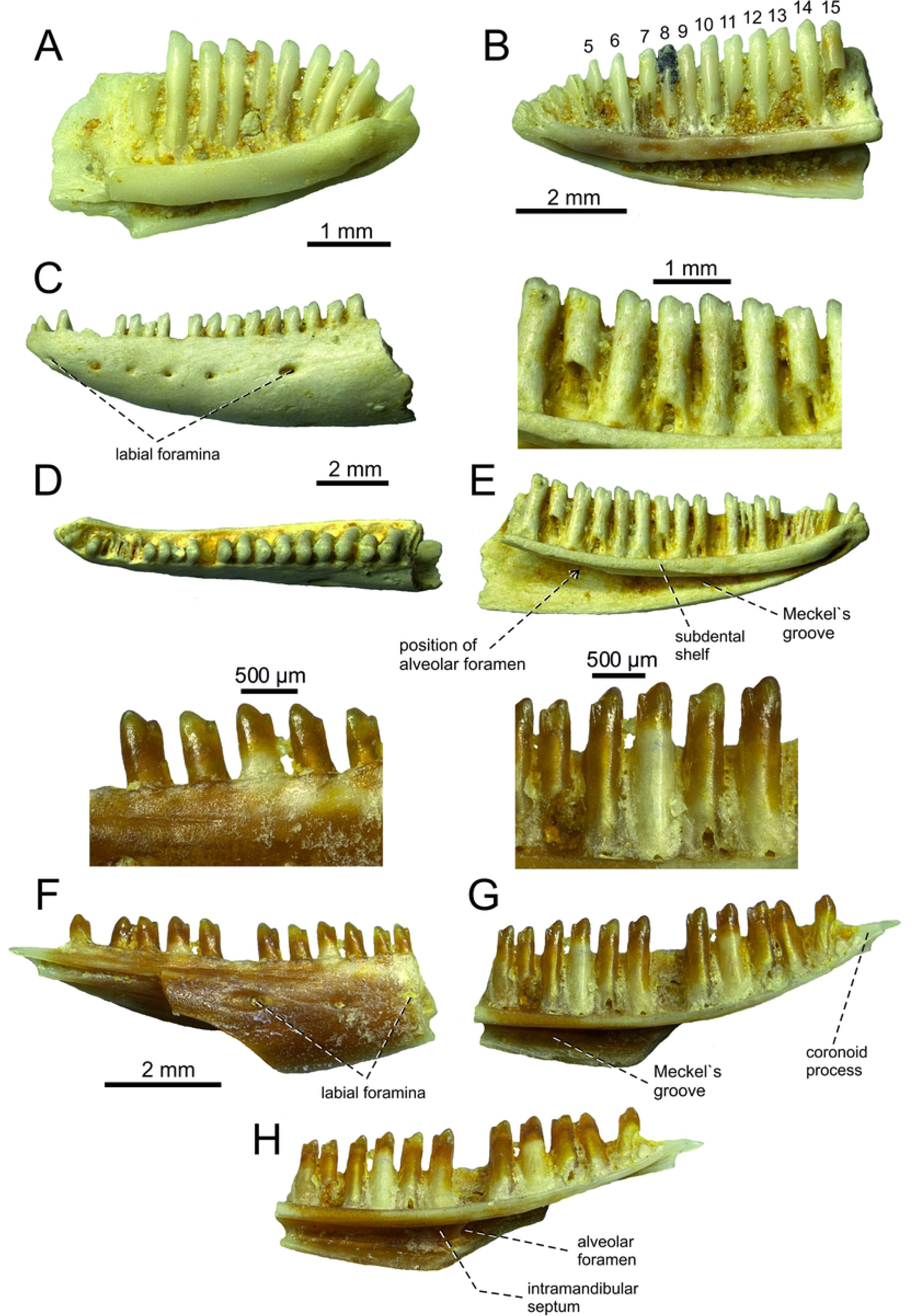
*Lacerta* cf. *trilineata* from the late Miocene of the locality Solnechnodolsk. Left dentary GIN 1145/294 in medial (A) aspect. Right dentary GIN1145/296 in medial (B) aspects. Left dentary GIN 1145/295 in lateral (C), dorsal (D) and medial (E) with detail of teeth. Right dentary GIN1145/297 in lateral (F) with detail of teeth, medial (G) with detail of teeth and ventromedial (H) aspects.

The otherwise smooth labial surface is pierced by six labial foramina, with the posteriomost foramen being the largest one. They are located in a single line located in the upper half of the dentary rather than being in the mid-region.

#### Dentition

The implantation is pleurodont. Teeth are tall, overarching the dental crest by the 1/3 of the tooth length. The tooth size slightly rises posteriorly. The anteriormost teeth are monocuspid, posteriorly bicuspid, starting from the 6th-7th tooth. There is a dominant cusp andanteriorly located accessory cusp. The small interdental gaps are present. Most of the tooth bases are pierced by resorption pits.

***Lacerta* group of ? *L. trilineata***

Fig. 8

**Fig 8.**
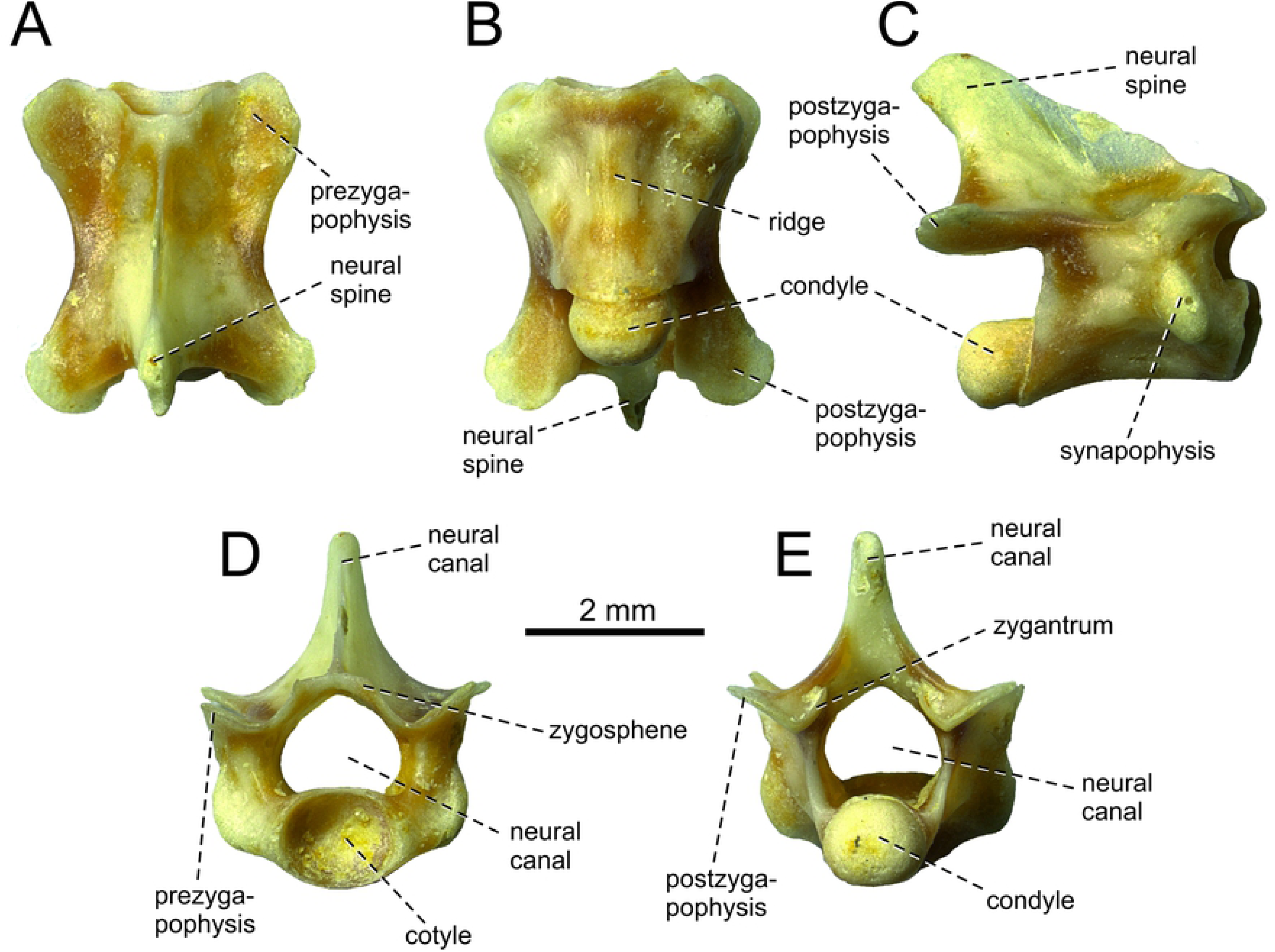
*Lacerta* group of ? *L. trilineata* from the late Miocene of the locality Solnechnodolsk. Dorsal vertebra GIN 1145/300 in dorsal (A), ventral (B), lateral (C), anterior (D) and posterior (E) aspects.

Material––Three vertebrae GIN 1145/300, 302, 303.

Locality and horizon––Solnechnodolsk (Stavropol Region, southwestern Russia); upper Miocene, MN 13.

### Osteological Description

#### Dorsal vertebra

Three dorsal vertebrae are preserved, of which the specimen GIN 1145/300 is almost completely preserved (Fig. 8).The neural spine is tall. In lateral aspect, the neural spine is posterodorsally oriented, trapezoidal - narrows posterodorsally. It slightly exceeds the condyle posteriorly. In dorsal aspect, the ascending region of the neural spine is lens shaped whereas the anteriorly located portion is thin. The interzygapophyseal constriction is only weakly developed. The prezygapophyses bear oval articulation surfaces. Postzygapophyses bear roughly triangular articulation surfaces, being more posteriorly oriented rather than laterally. Zygosphene-zygantrum accessory intervertebral articulations are present, but only incipient, weakly developed as small facets located at the base of the neural arch and continuous with pre- or - postzygapophyseal articulations (character 468, state 2 in [45]). The neural canal is large, pentagonal in shape. The synapophysis is present in the anterior half of the vertebrae, being large and ovoid shaped. The posterior centrosynapophyseal lamina (sensu Tschopp [46]) is present posteriorly to the synapophyses. The cotyle and condyle are rounded. In ventral aspect, the centrum bears a low ridge in its mid-region. In lateral aspect, the ventral margin of the centrum is concave.

#### Remarks

The dorsal vertebrae of green lizards have tendency to uniform morphology, which encourages caution in assigning isolated vertebrae to particular species. The dorsal vertebrae from Solnechnodolsk shows a high similarity to that of *L. trilineata*, rather than to e.g. *L. strigata*. The posterior centrosynapophyseal lamina is absent in *L. strigata* and the synapophysis here is small (for vertebra of *L. trilineata* and *L. strigata*, see [46]). There are no characters that would prevent this vertebra from being assigned to *L. trilineata*, but strong support for such statement is absent. For this reason, we allocate this material here only potentionally to the group of *L. trilineata*.

#### Discussion and comparison

All fossil bones described here can be allocated to Lacertidae without doubt. They are assigned to a single species (except of vertebrae, see above) on the basis of dentition, sculpture of osteodermal shields and great similarity with the morphology presented by the extant green lizards (see below). Moreover they are comparable in size and come from the single site. Green lizards form a uniform group of lacertids. Their fossil record is rather poor (see Introduction), what is caused by two factors: (a) the low number of known fossil elements with a proper diagnostic features; (b) the lack of the comparative studies focused on the detailed comparison of osteological elements in this group. For a proper identification of the Solnechnodolsk fossils, it was necessary to present such a study here. Although *Lacerta agilis* is traditionally called the sand lizard, it has been shown that this taxon branches within the green lizards [16, 26]. For this reason, it is included here and the complete list of green lizards studied here includes *L. agilis, L. viridis, L. bilineata, L. pamphylica, L. strigata, L. schreiberi, L. trilineata* and *L. media* (see Figs. 9–26). *Timon lepidus* was used for outgroup comparison (Figs. 27, 28). But because *T. lepidus* has a lot of unique features among extant lacertids (see below) and, moreover, to record the distribution of some character states, we also used *P. muralis* (Fig. 29), *Zootoca vivipara* (Fig. 30), *Takydromus sexlineatus* (Fig. 31A-F), *Meroles ctenodactylus* (Fig. 31G-I), *Psammodromus algirus* (Fig. 32) and *Gallotia stehlini* (Fig. 34).

**Fig 9.**
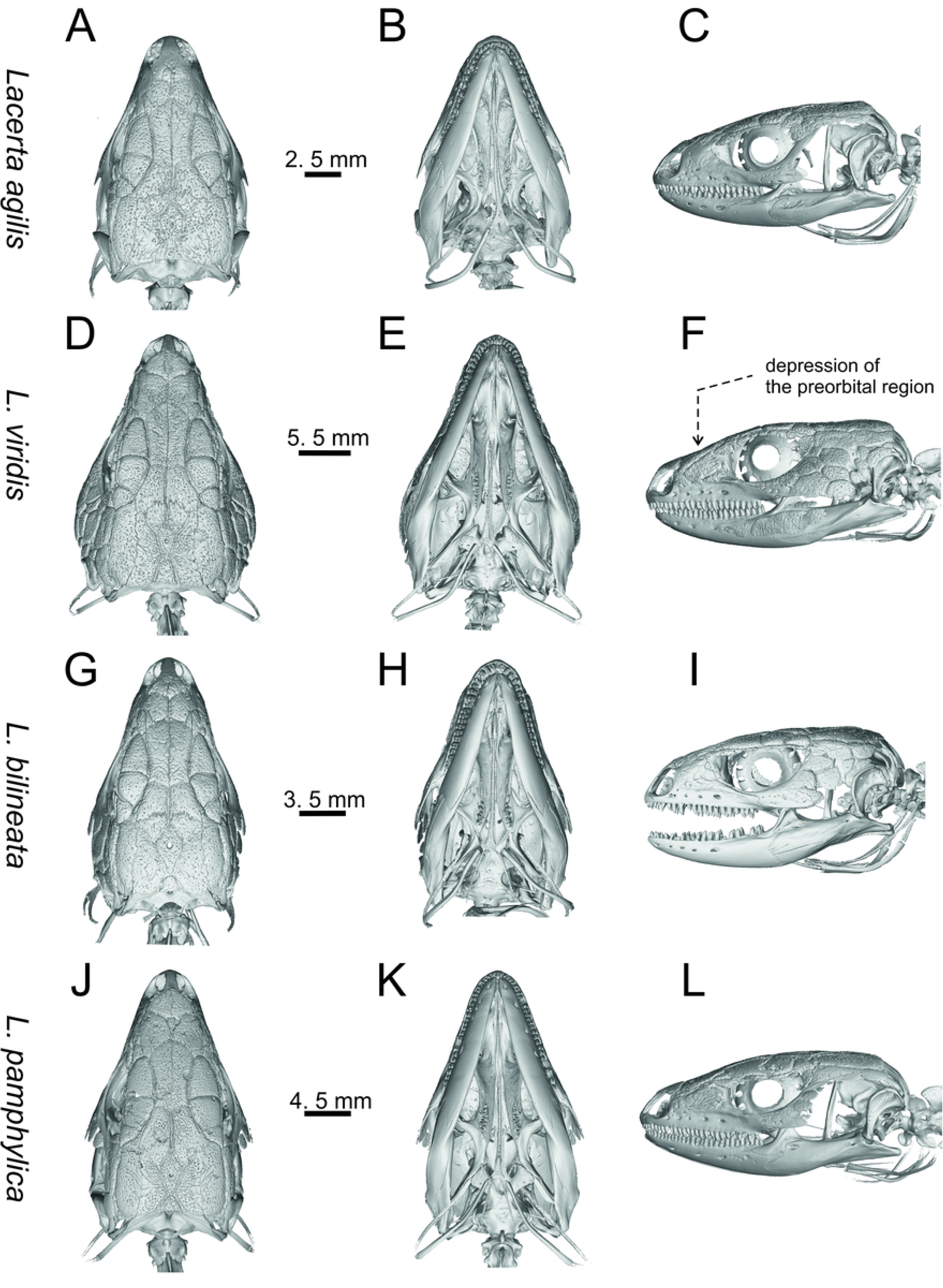
Skulls of extant green lizards *Lacerta agilis, L. viridis, L bilineata* and *L. pamphylica*. In dorsal (A, D, G, J), ventral (B, E, H, K) and lateral (C, F, I, L) aspects (continued).

**Fig 10.**
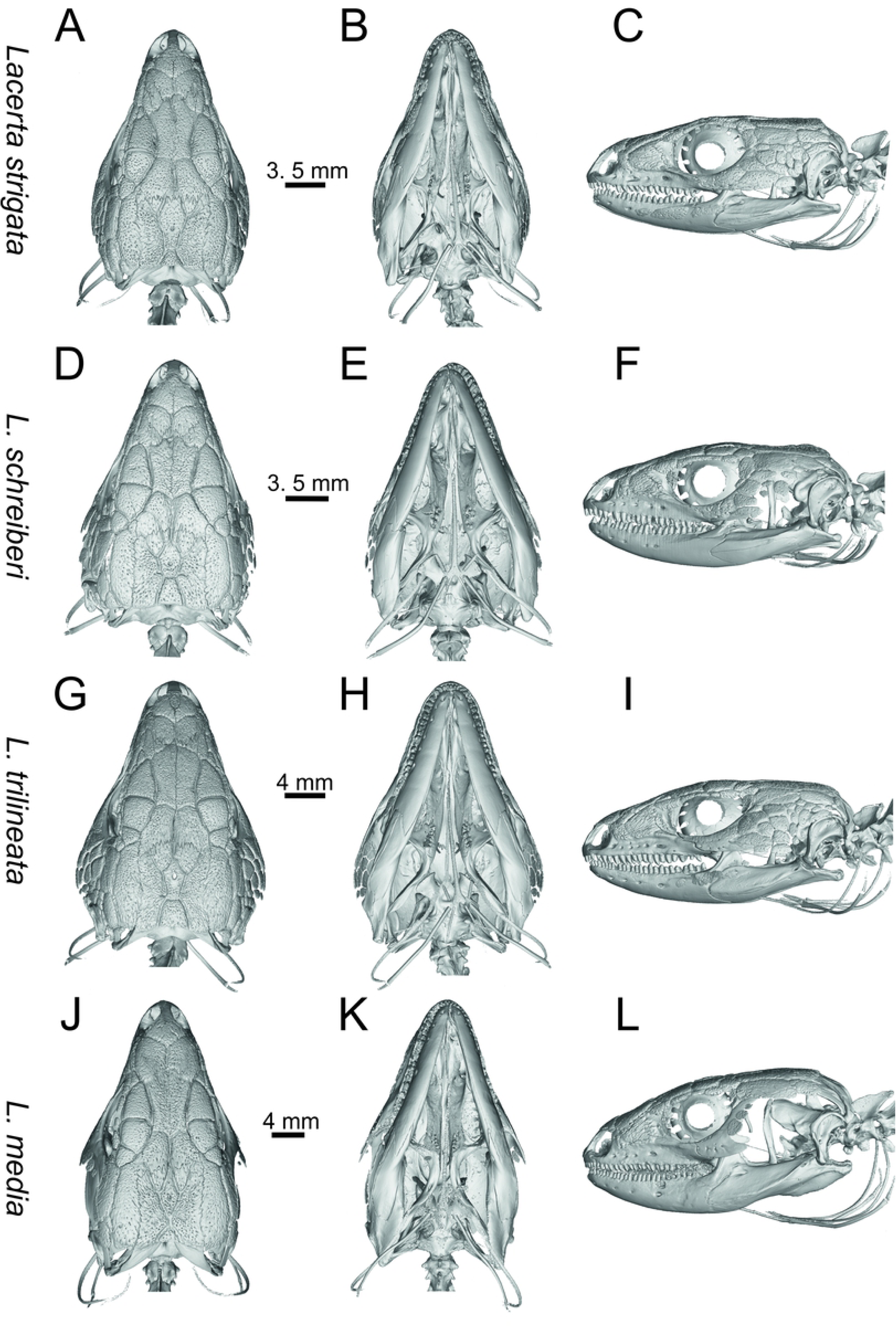
Skulls of extant green lizards *Lacerta strigata, L. schreiberi, L trilineata* and *L. media*. In dorsal (A, D, G, J), ventral (B, E, H, K) and lateral (C, F, I, L) aspects.

**Fig 11.**
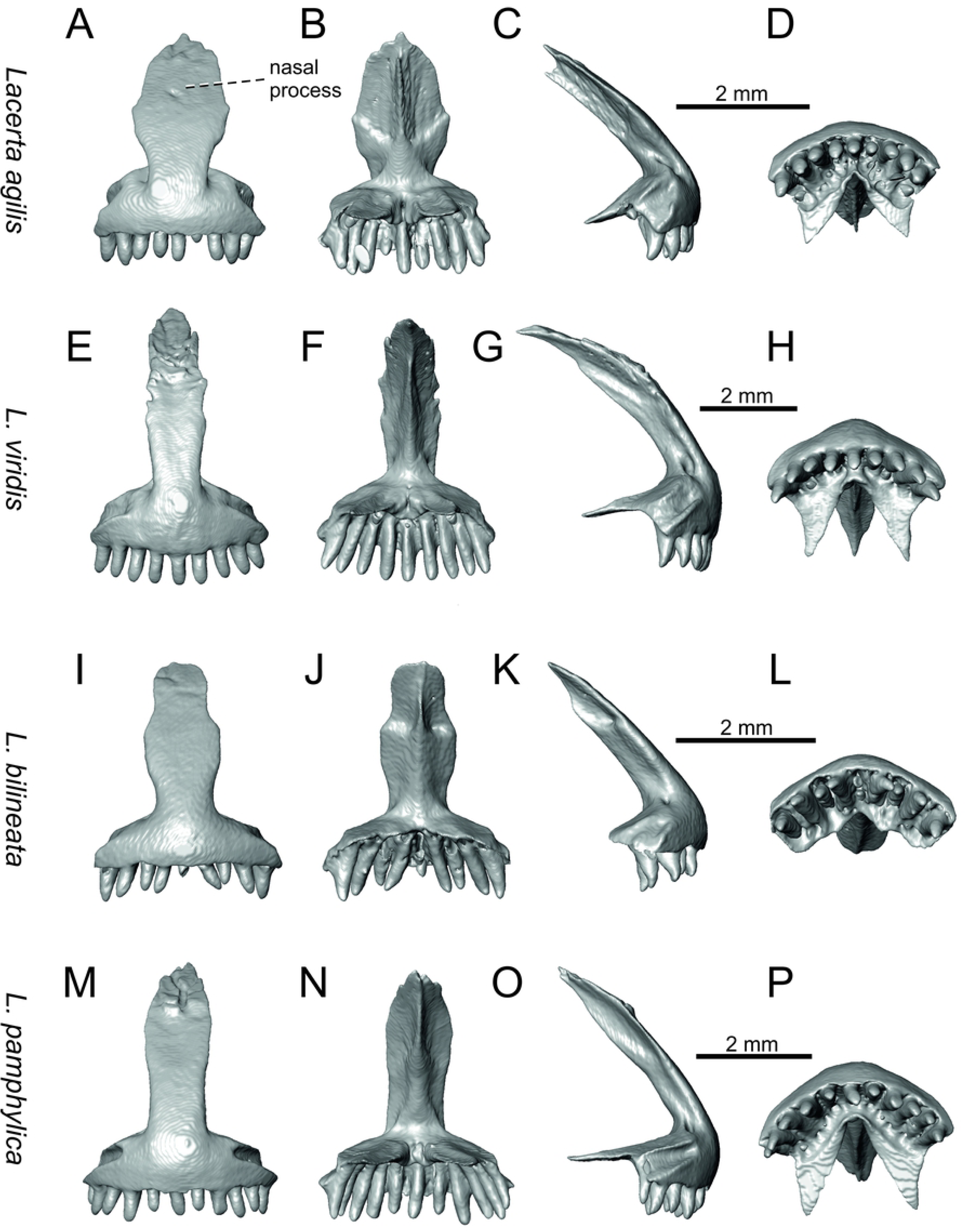
Premaxillae of extant green lizards *Lacerta agilis, L. viridis, L bilineata* and *L. pamphylica*. In anterior (A, E, I, M), posterior (B, F, J, N), lateral (C, G, K, O) and ventral (D, H, L, P) aspects (continued).

**Fig 12.**
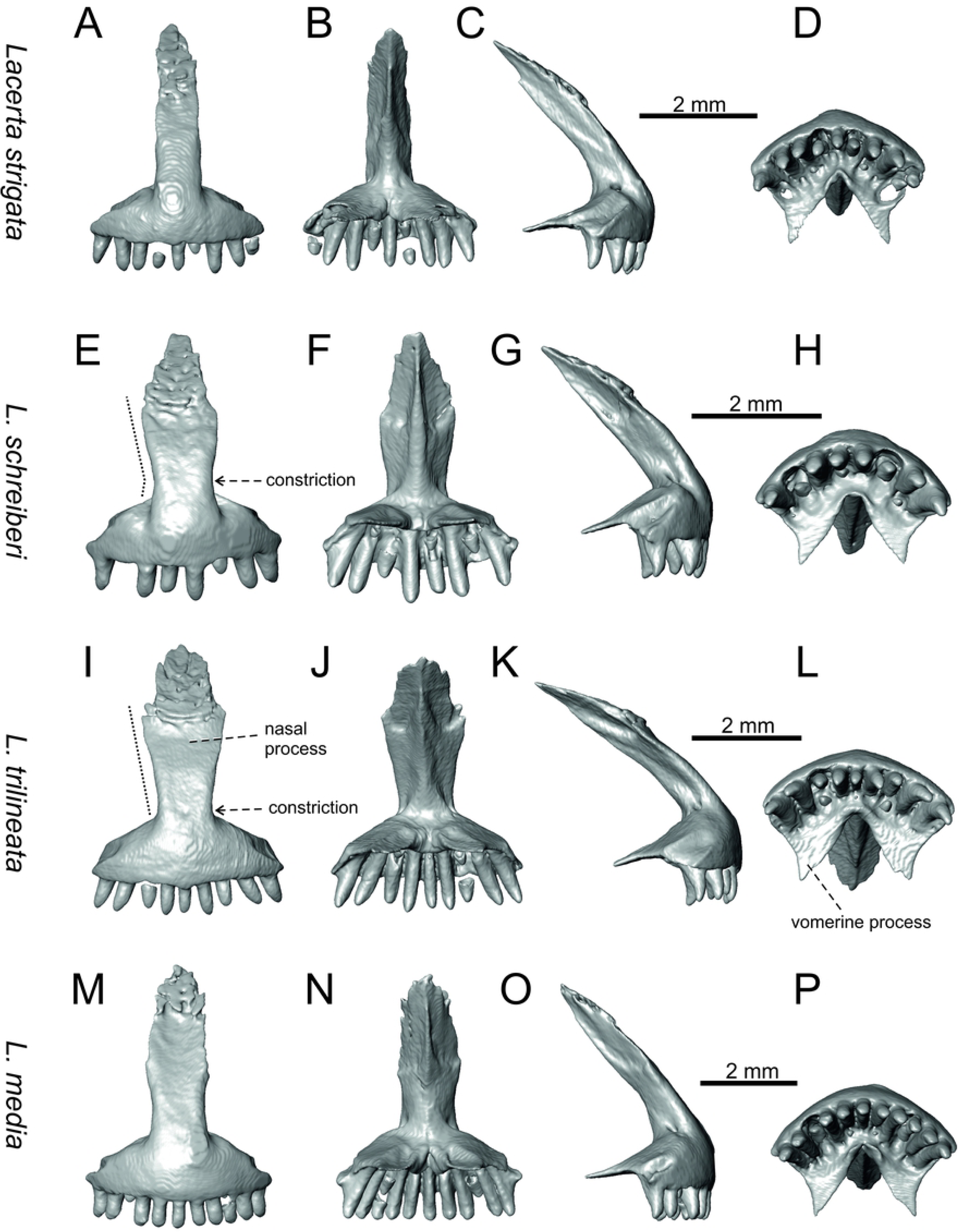
Premaxillae of extant green lizards *Lacerta strigata, L. schreiberi, L trilineata* and *L. media*. In anterior (A, E, I, M), posterior (B, F, J, N), lateral (C, G, K, O) and ventral (D, H, L, P) aspects. The dotted lines indicate a lateral margin of the nasal process and a position of constriction in *L. schreiberi* and *L. trilineata*.

**Fig 13.**
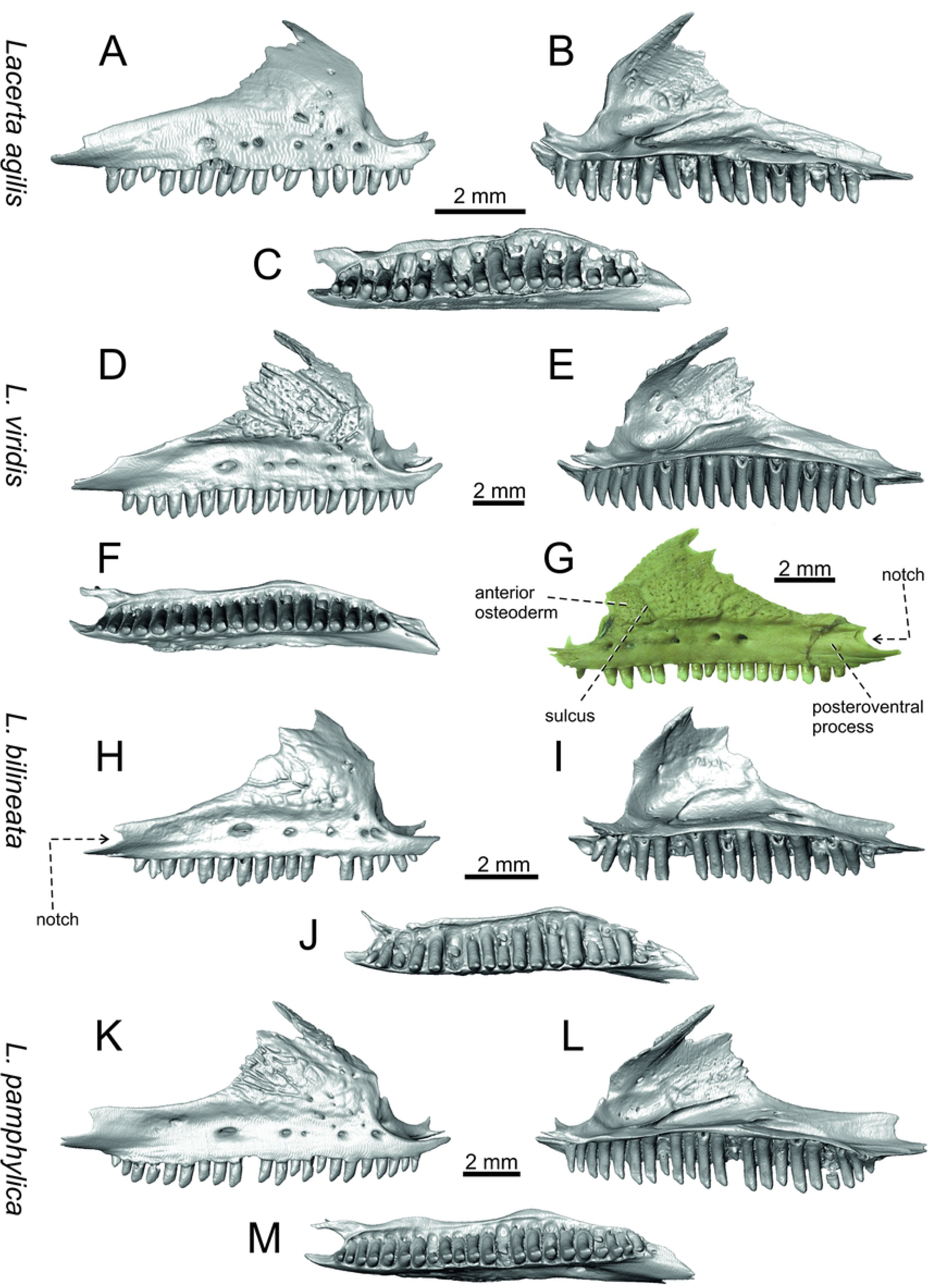
Maxillae of extant green lizards *Lacerta agilis, L. viridis, L bilineata* and *L. pamphylica*. In lateral (A, D, G, H, K), medial (B, E, I, L) and ventral (C, F, J, M) aspects (continued).

**Fig 14.**
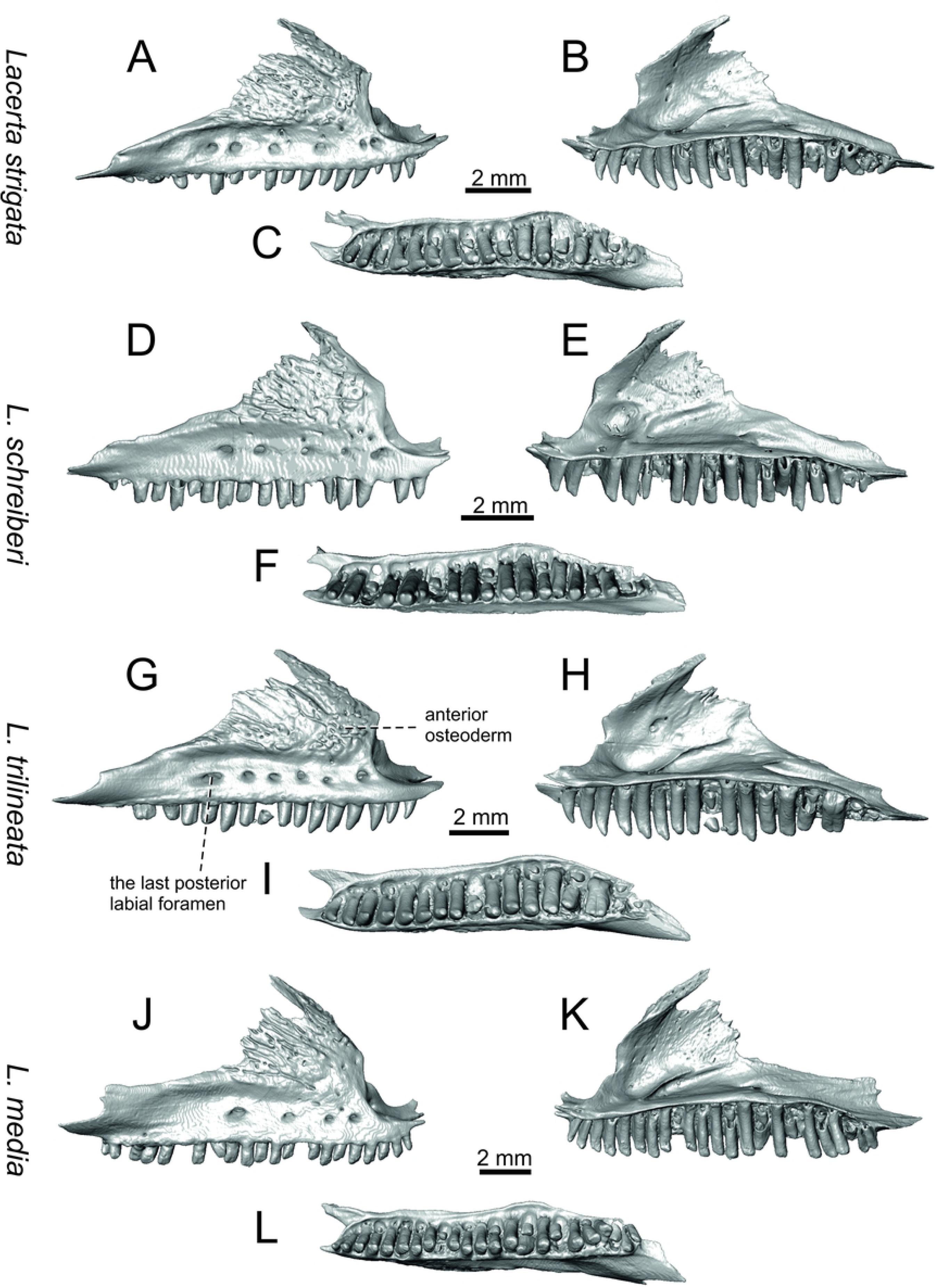
Maxillae of extant green lizards *Lacerta strigata, L. schreiberi, L trilineata* and *L. media*. In lateral (A, D, G, J), medial (B, E, H, K) and ventral (C, F, I, L) aspects (continued).

**Fig 15.**
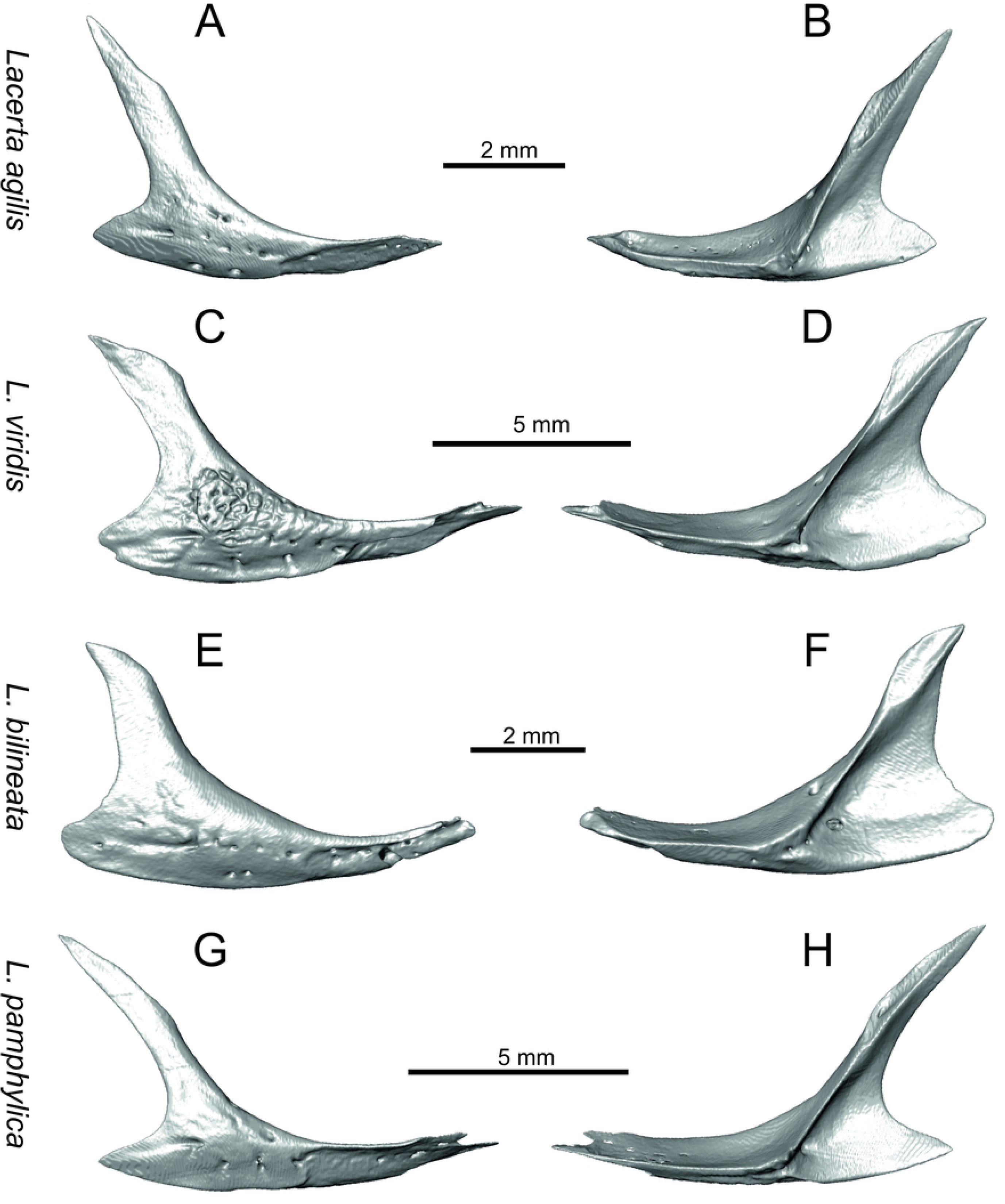
Jugals of extant green lizards *Lacerta agilis, L. viridis, L bilineata* and *L. pamphylica*. In lateral (A, C, E, G) and medial (B, D, F, H) aspects (continued).

**Fig 16.**
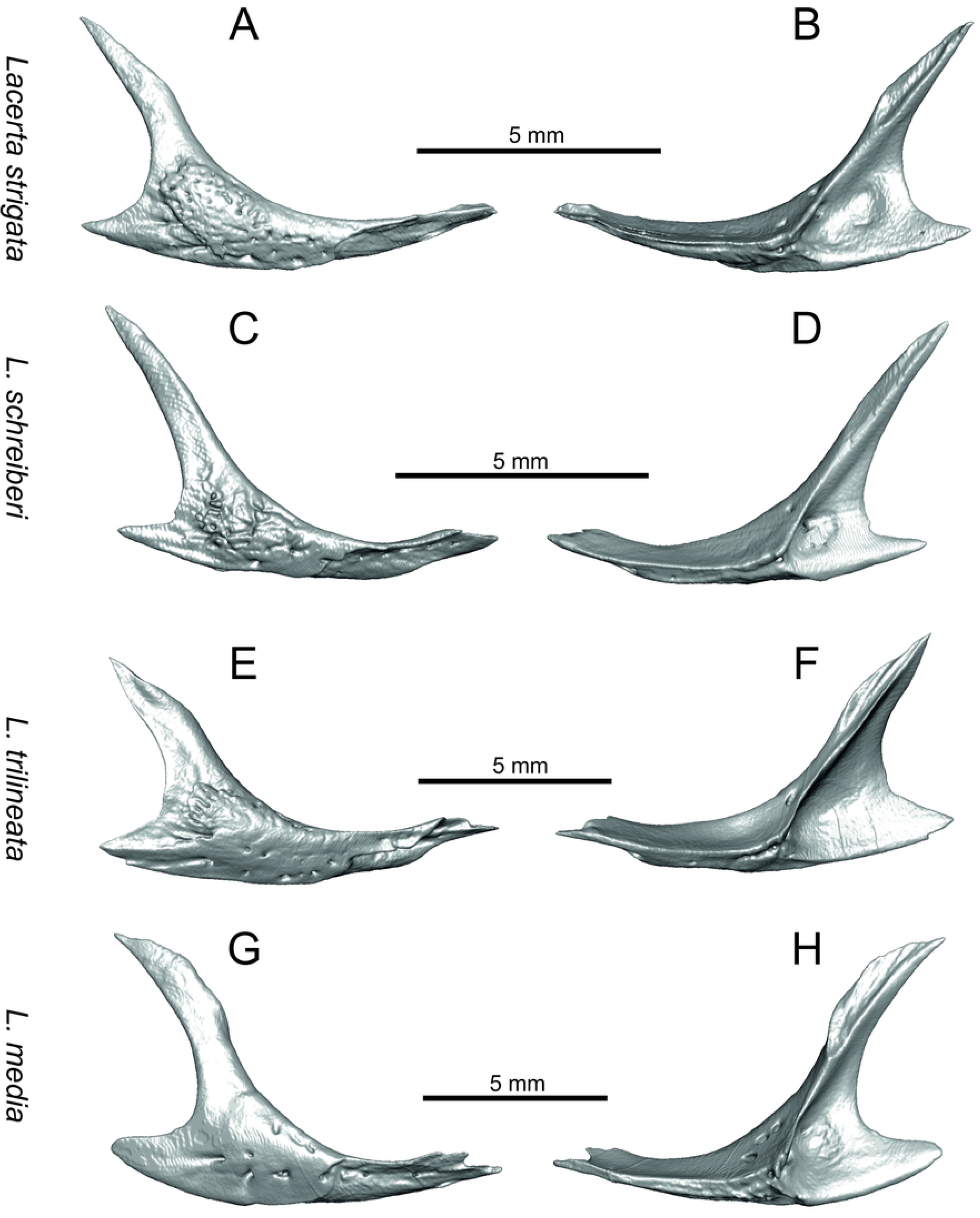
Jugals of extant green lizards *Lacerta strigata, L. schreiberi, L trilineata* and *L. media*. In lateral (A, C, E, G) and medial (B, D, F, H) aspects.

**Fig 17.**
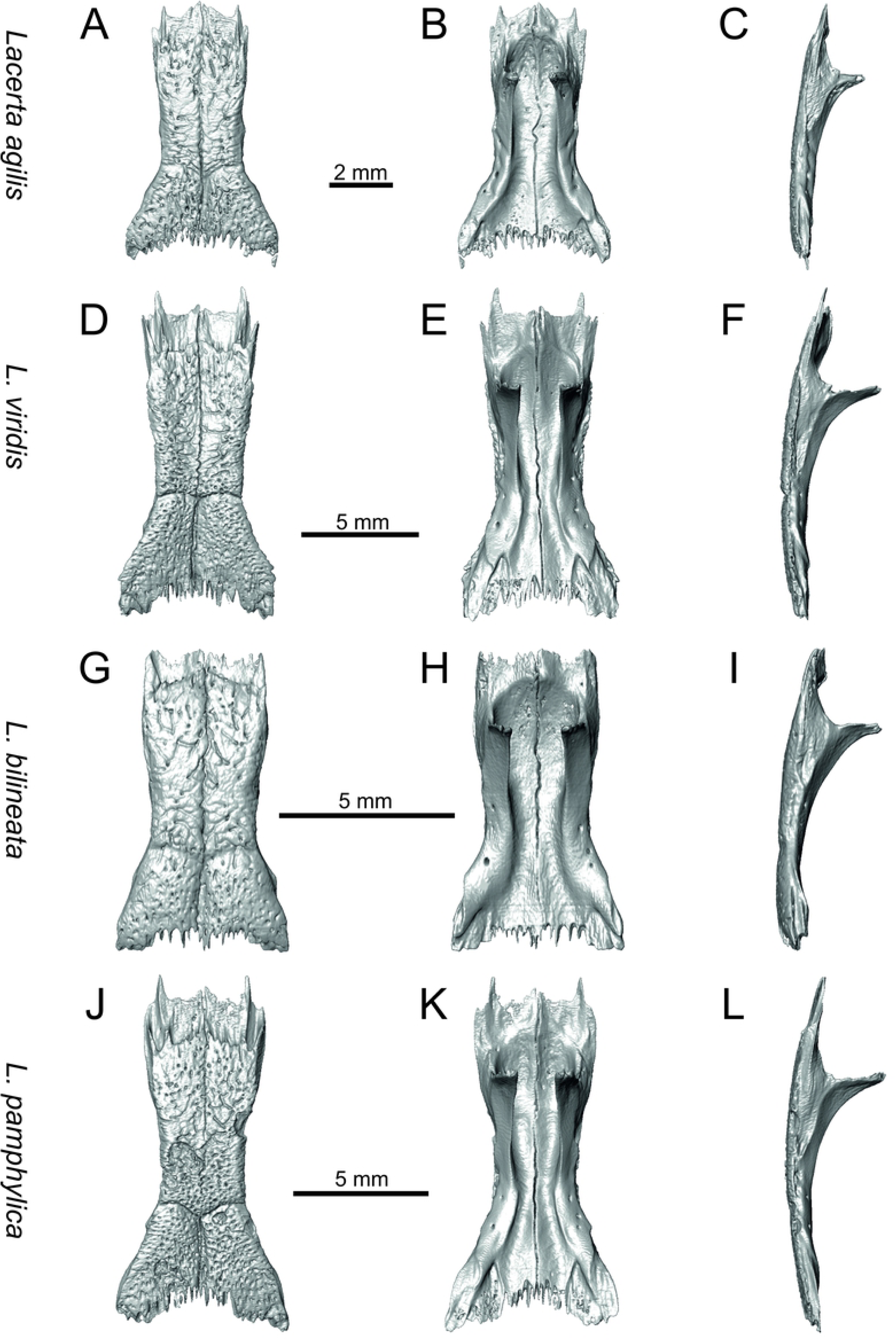
Frontals of extant green lizards *Lacerta agilis, L. viridis, L bilineata* and *L. pamphylica*. In dorsal (A, D, G, J), ventral (B, E, H, K) and lateral (C, F, I, L) aspects (continued).

**Fig 18.**
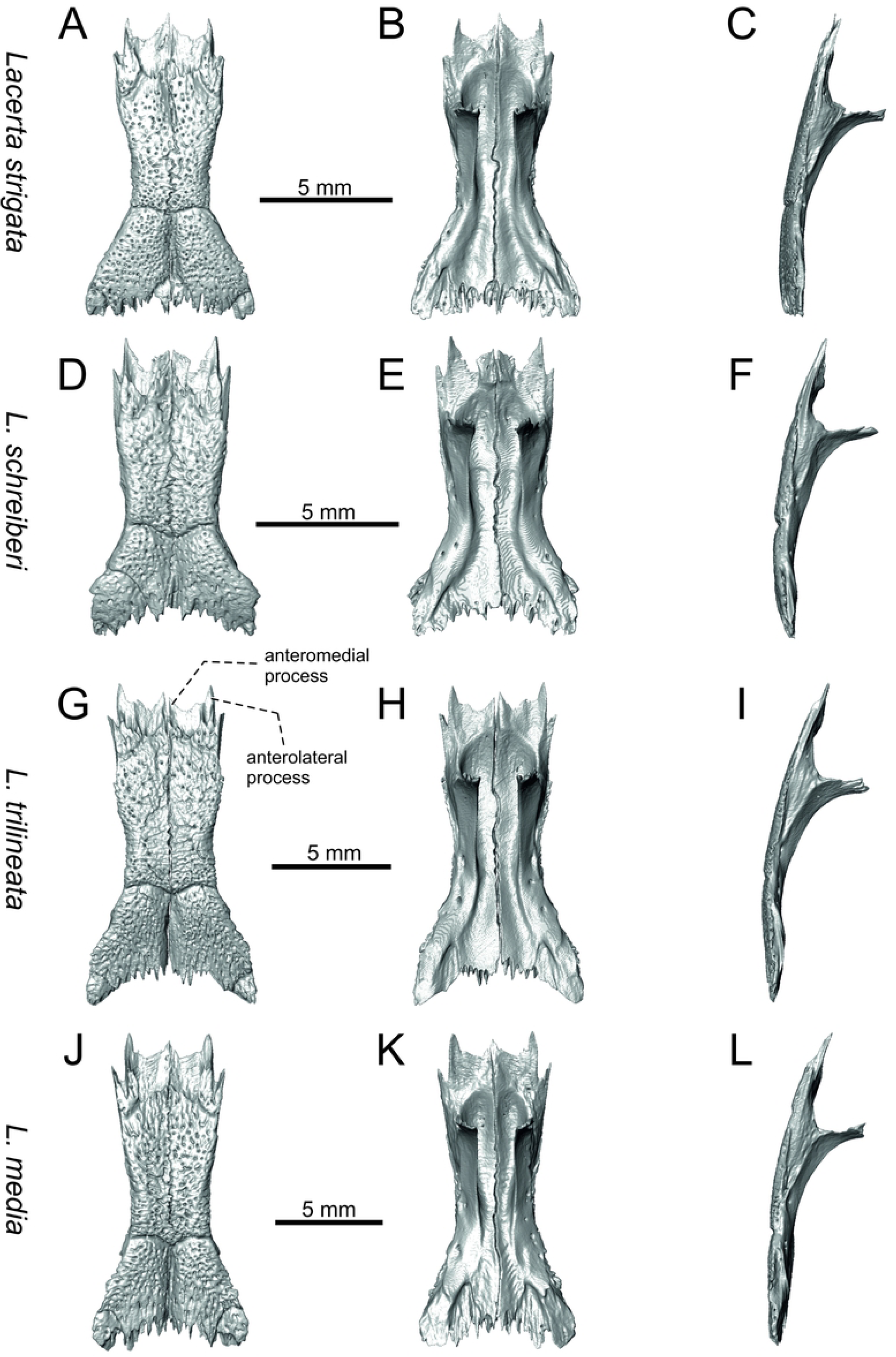
Frontals of extant green lizards *Lacerta strigata, L. schreiberi, L trilineata* and *L. media*. In dorsal (A, D, G, J), ventral (B, E, H, K) and lateral (C, F, I, L) aspects.

**Fig 19.**
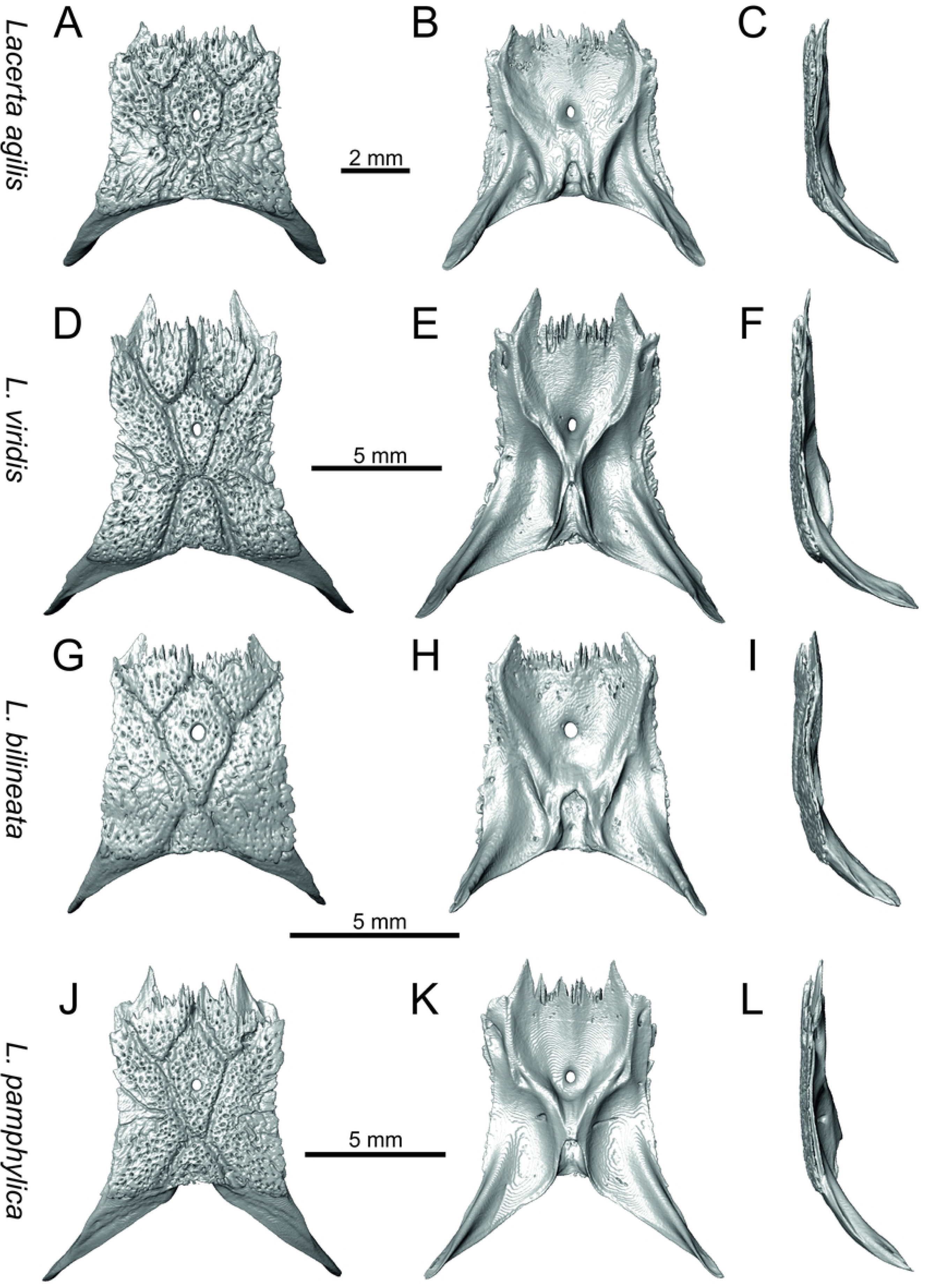
Parietals of extant green lizards *Lacerta agilis, L. viridis, L bilineata* and *L. pamphylica*. In dorsal (A, D, G, J), ventral (B, E, H, K) and lateral (C, F, I, L) aspects (continued).

**Fig 20.**
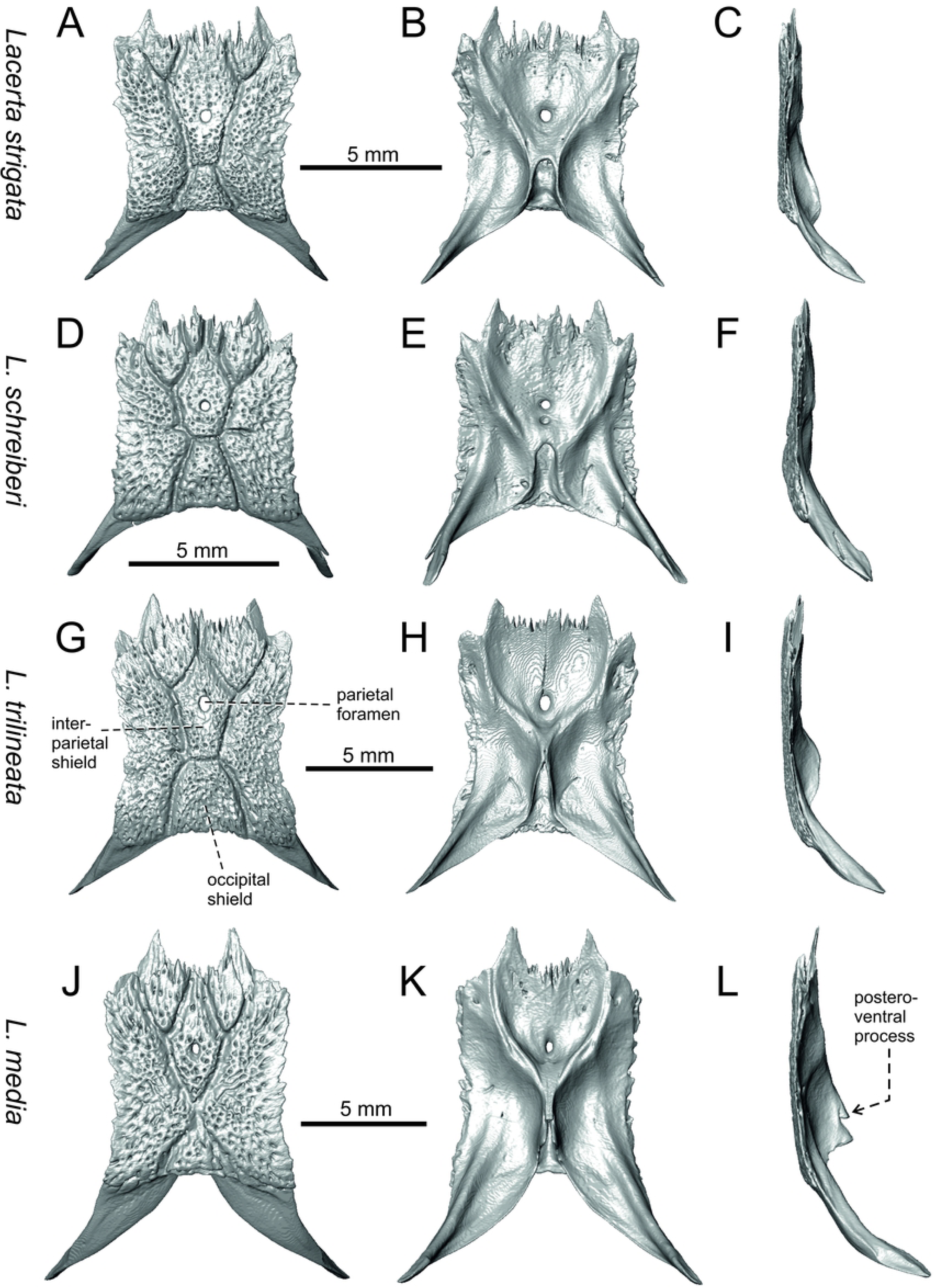
Parietals of extant green lizards *Lacerta strigata, L. schreiberi, L trilineata* and *L. media*. In dorsal (A, D, G, J), ventral (B, E, H, K) and lateral (C, F, I, L) aspects.

**Fig 21.**
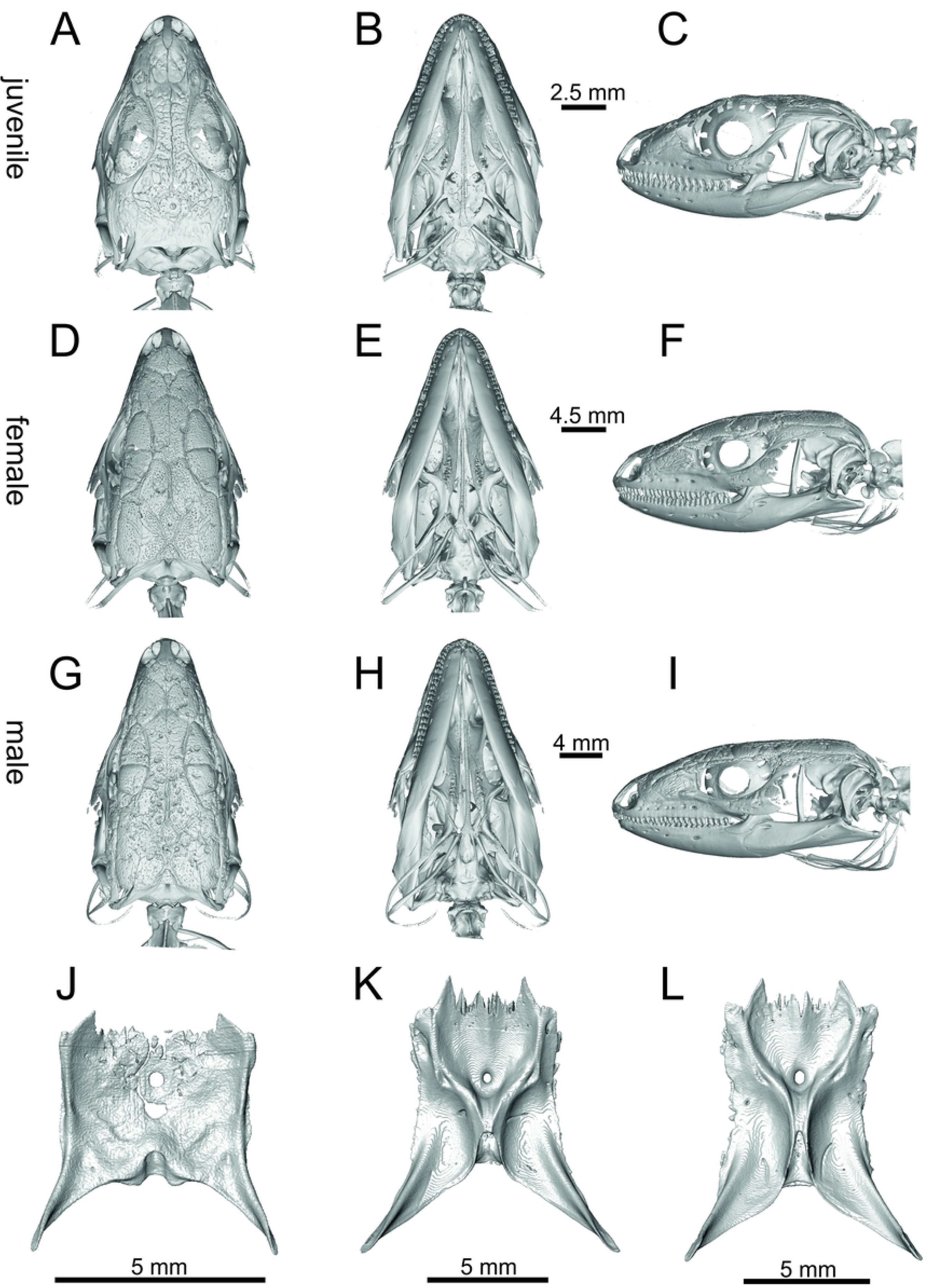
Ontogenetic and sexual variety of *Lacerta pamphylica*. (juvenile A-C, J; female D-F, K and male G-I, L): skulls in dorsal (A, D, G), ventral (B, E, H) and lateral (C, F, I) apsects. Parietals in ventral (J, K, L) aspects.

**Fig 22.**
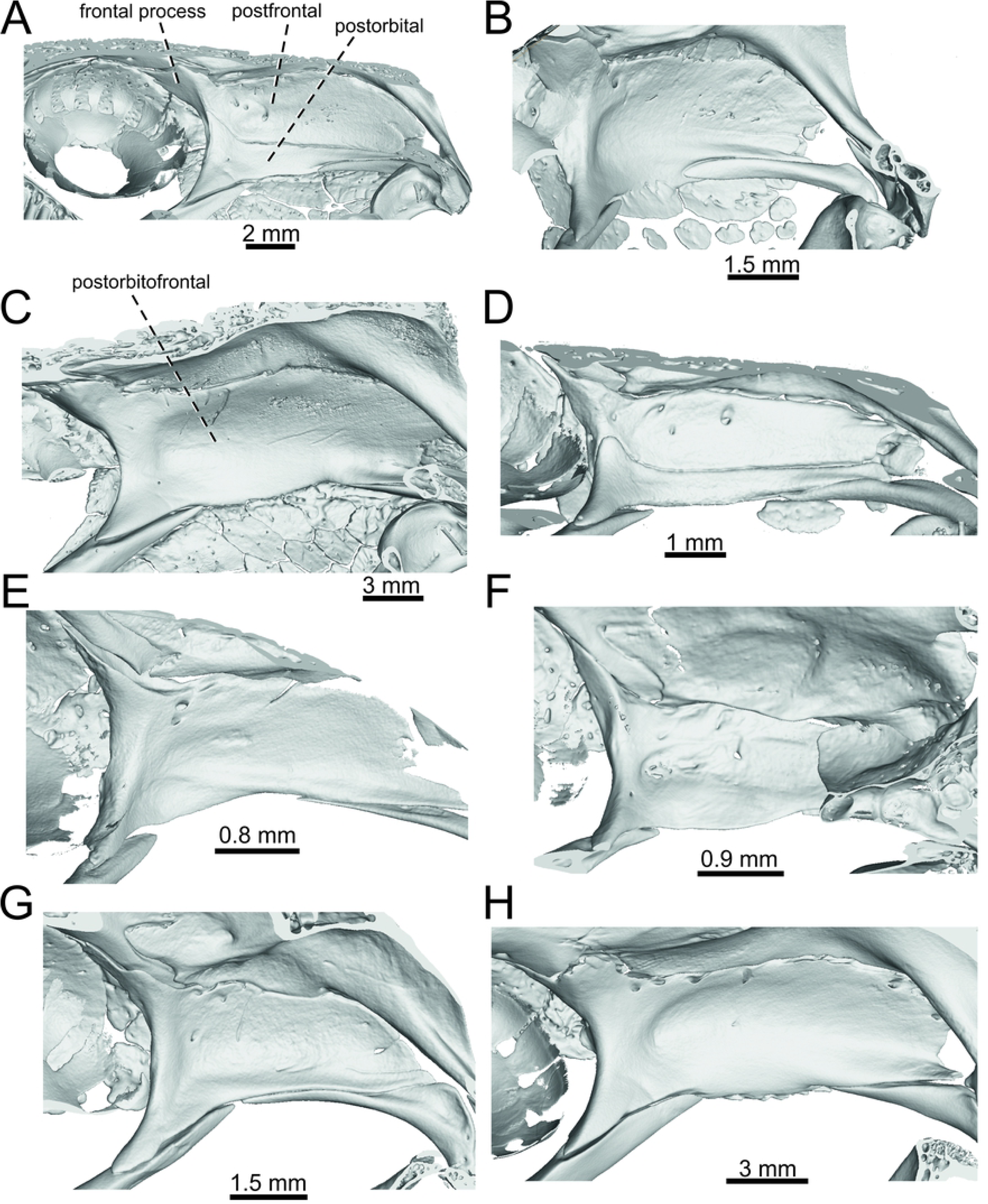
Postfrontals + postorbitals located in the skulls in internal aspects in *Lacerta viridis* (A), *L. schreiberi* (B), *Timon lepidus* (C), *Podarcis muralis* (D), *Zootoca vivipara* (E), *Takydromus sexlineatus* (F), *Meroles ctenodactylus* (G), *Gallotia stehlini* (H).

**Fig 23.**
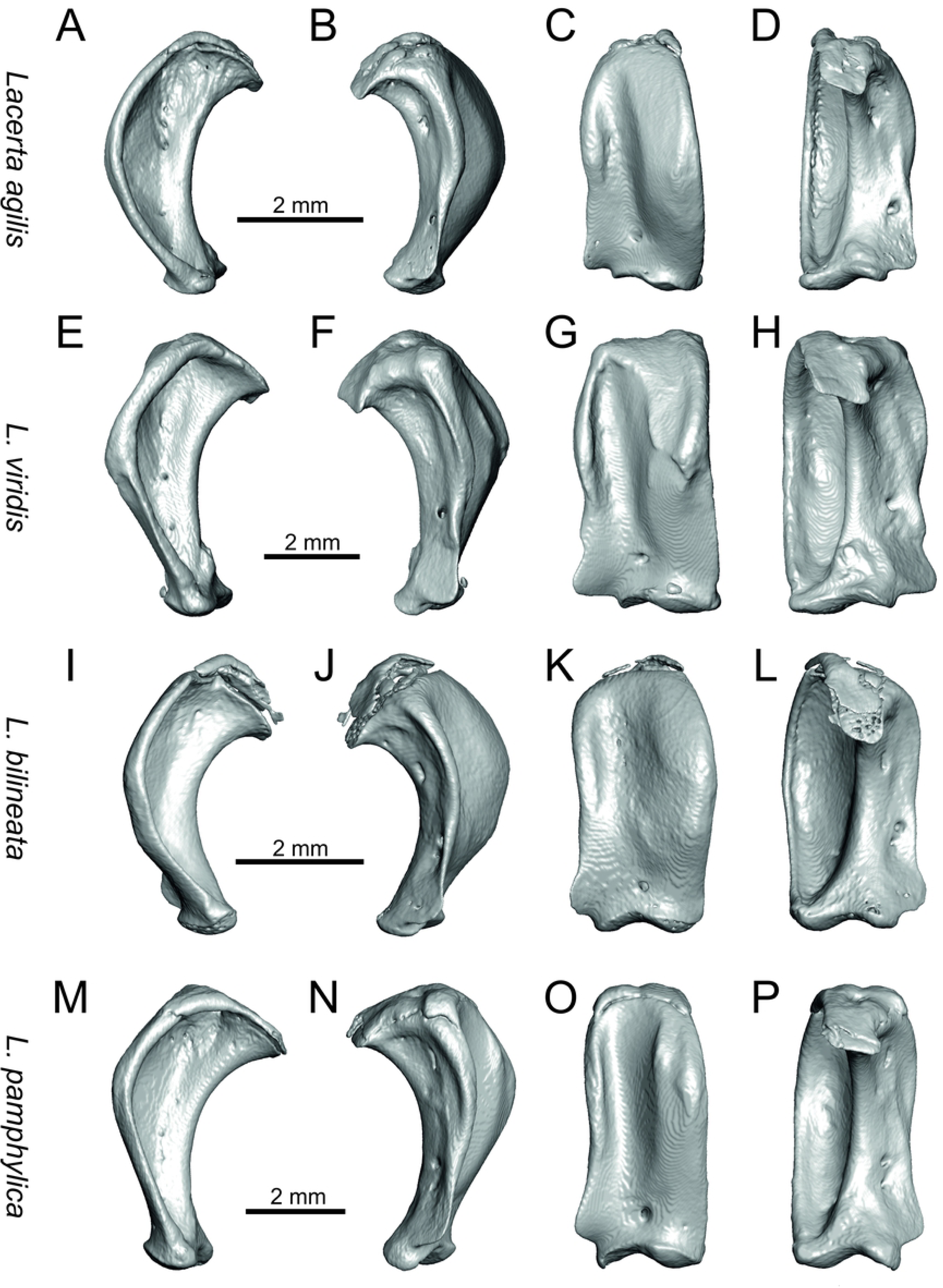
Quadrates of extant green lizards *Lacerta agilis, L. viridis, L bilineata* and *L. pamphylica*. In lateral (A, E, I, M), medial (B, F, J, N), anterior (C, G, K, O) and posterior (D, H, L, P) aspects (continued).

**Fig 24.**
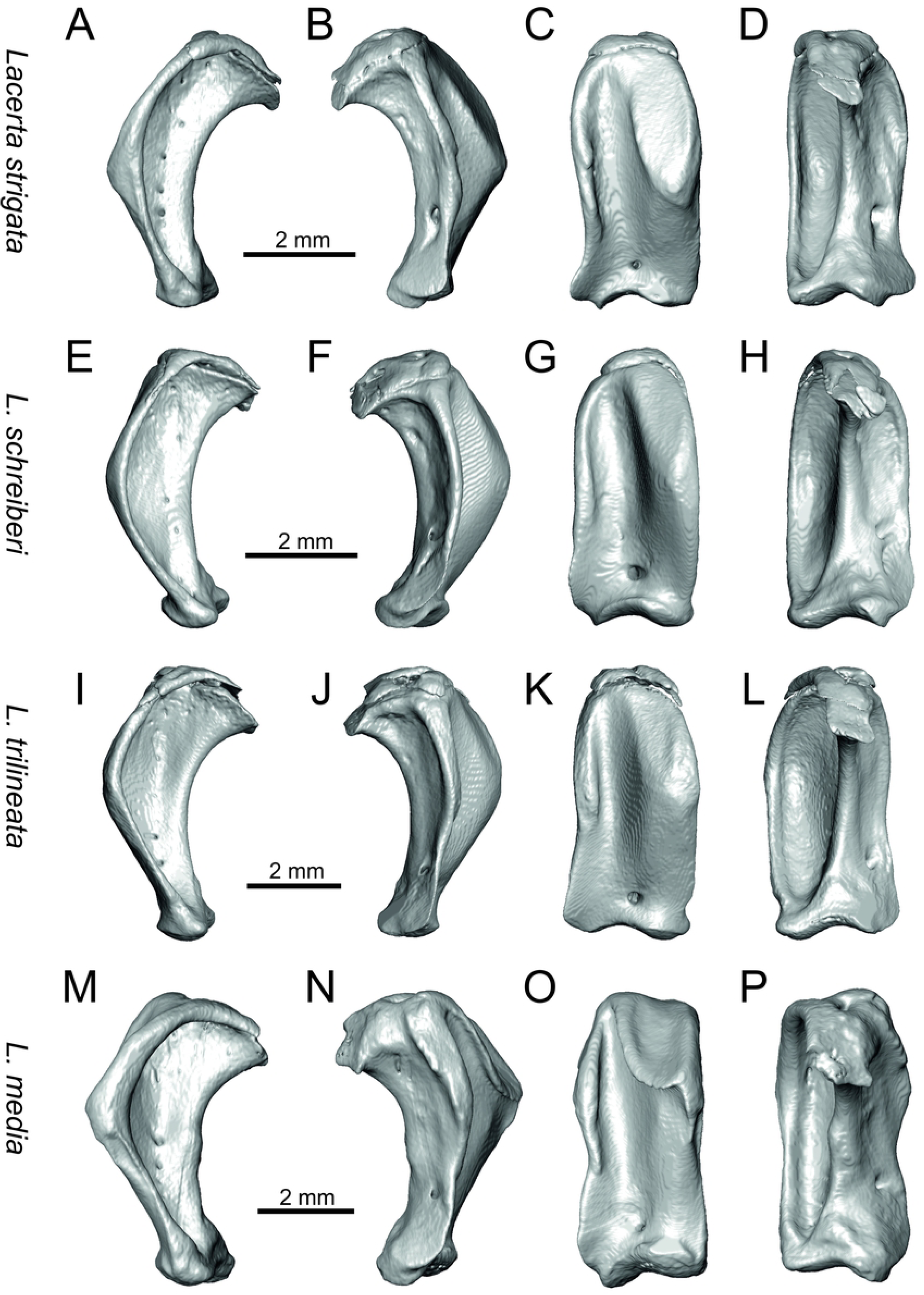
Quadrates of extant green lizards *Lacerta strigata, L. schreiberi, L trilineata* and *L. media*. In lateral (A, E, I, M), medial (B, F, J, N), anterior (C, G, K, O) and posterior (D, H, L, P) aspects.

**Fig 25.**
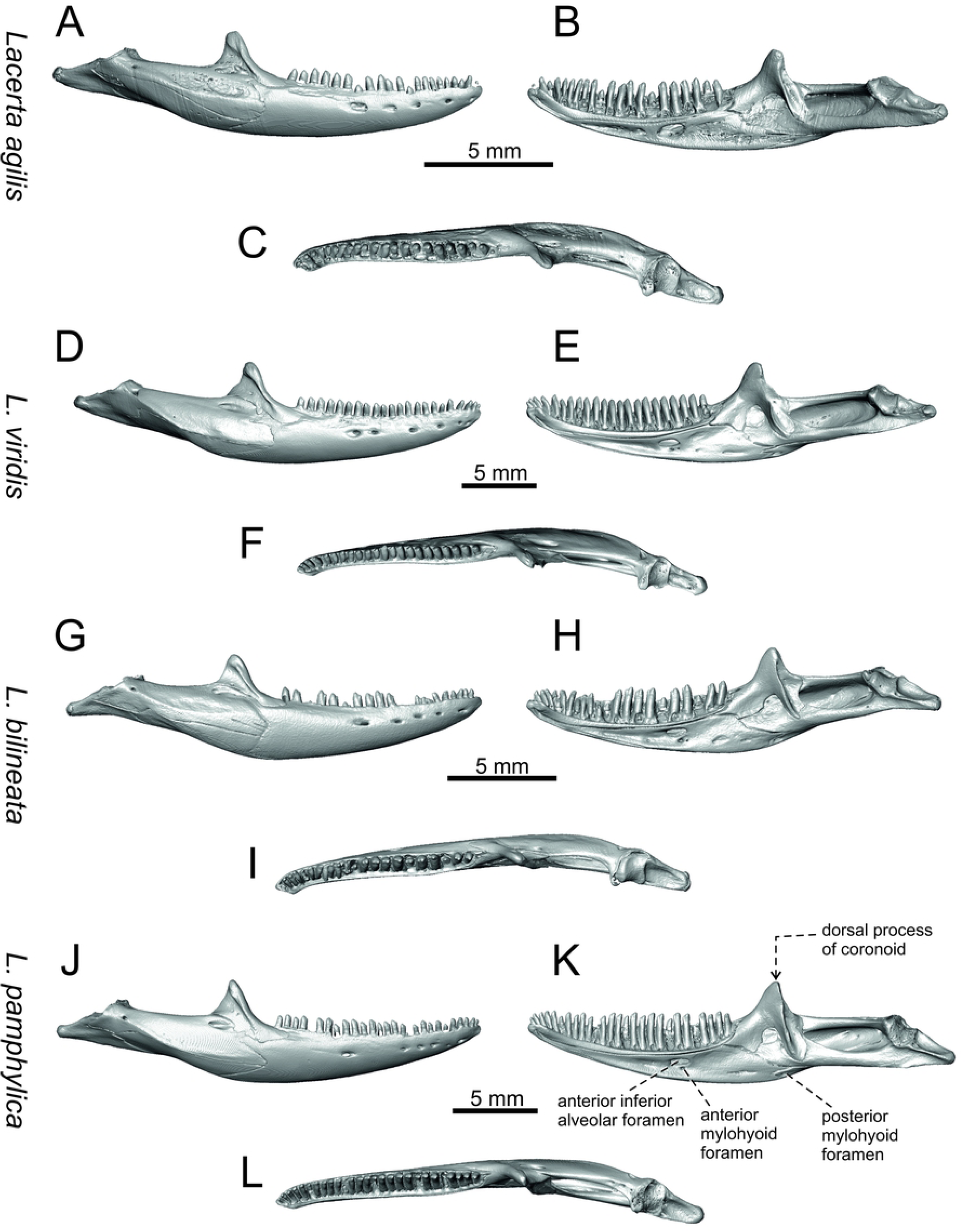
Mandibles of extant green lizards *Lacerta agilis, L. viridis, L. bilineata* and *L. pamphylica*. In lateral (A, D, G, J), medial (B, E, H, K) and dorsal (C, F, I, L) aspects (continued).

**Fig 26.**
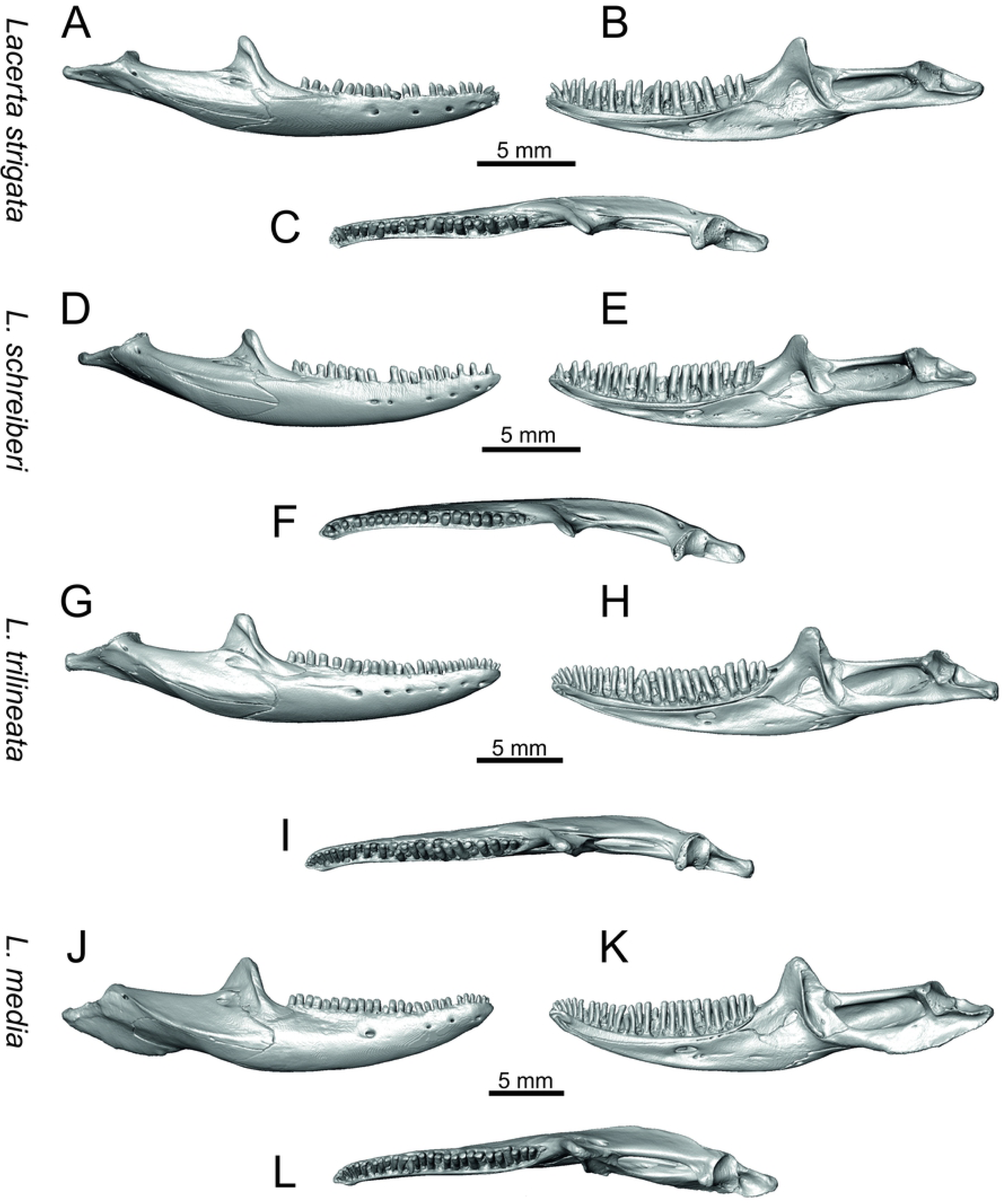
Mandibles of extant green lizards *Lacerta strigata, L. schreiberi, L trilineata* and *L. media*. In lateral (A, D, G, J), medial (B, E, H, K) and ventral (C, F, I, L) aspects (continued).

**Fig 27.**
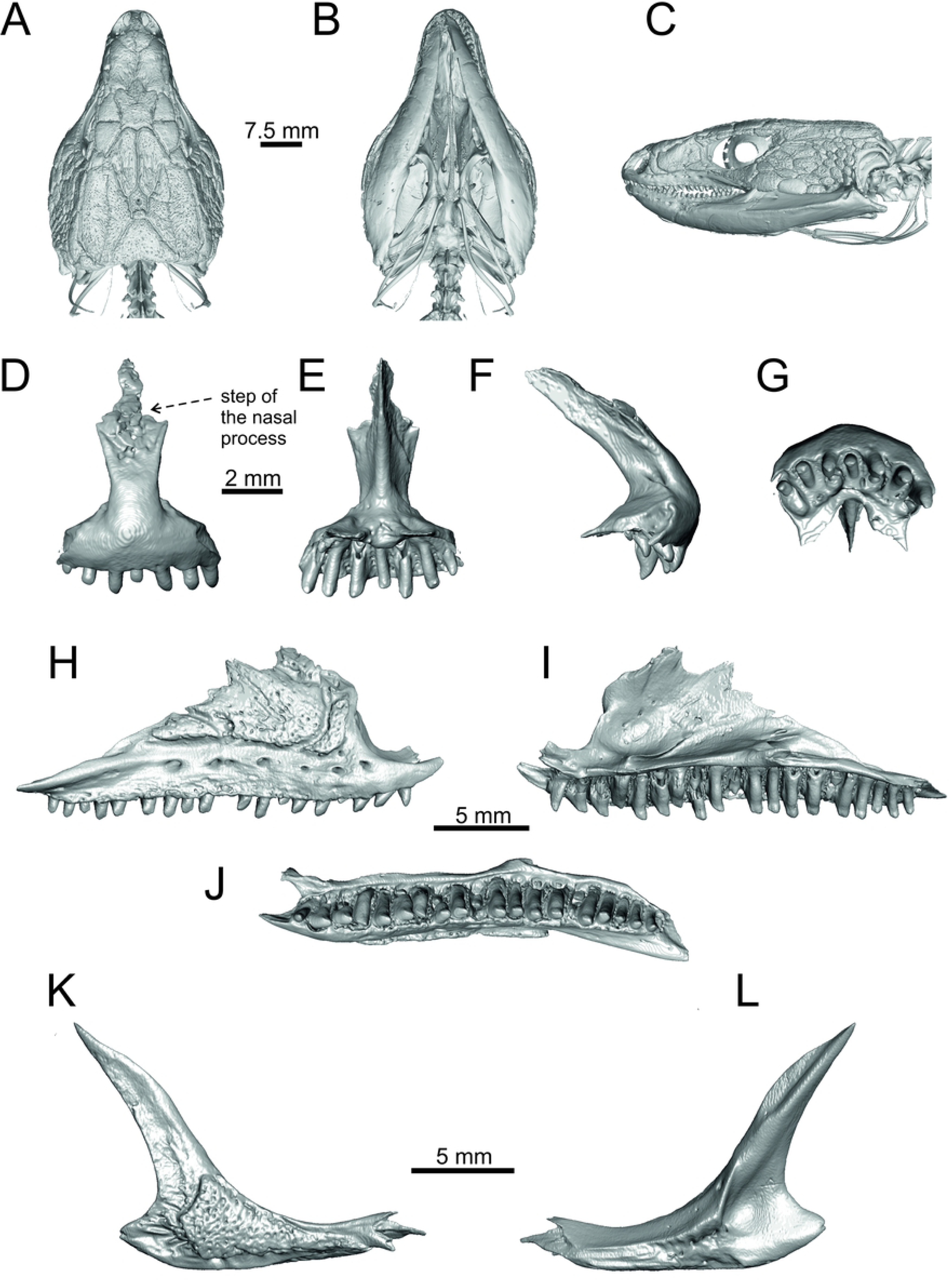
Skull and selected cranial elements of extant *Timon lepidus*. Skull in dorsal (A), ventral (B) and lateral (C) aspects. Premaxilla in anterior (D), posterior (E), lateral (F) and ventral (G) aspects. Maxilla in lateral (H), medial (I) and ventral (J) aspects. Jugal in lateral (K) and medial (L) aspects (continued).

**Fig 28.**
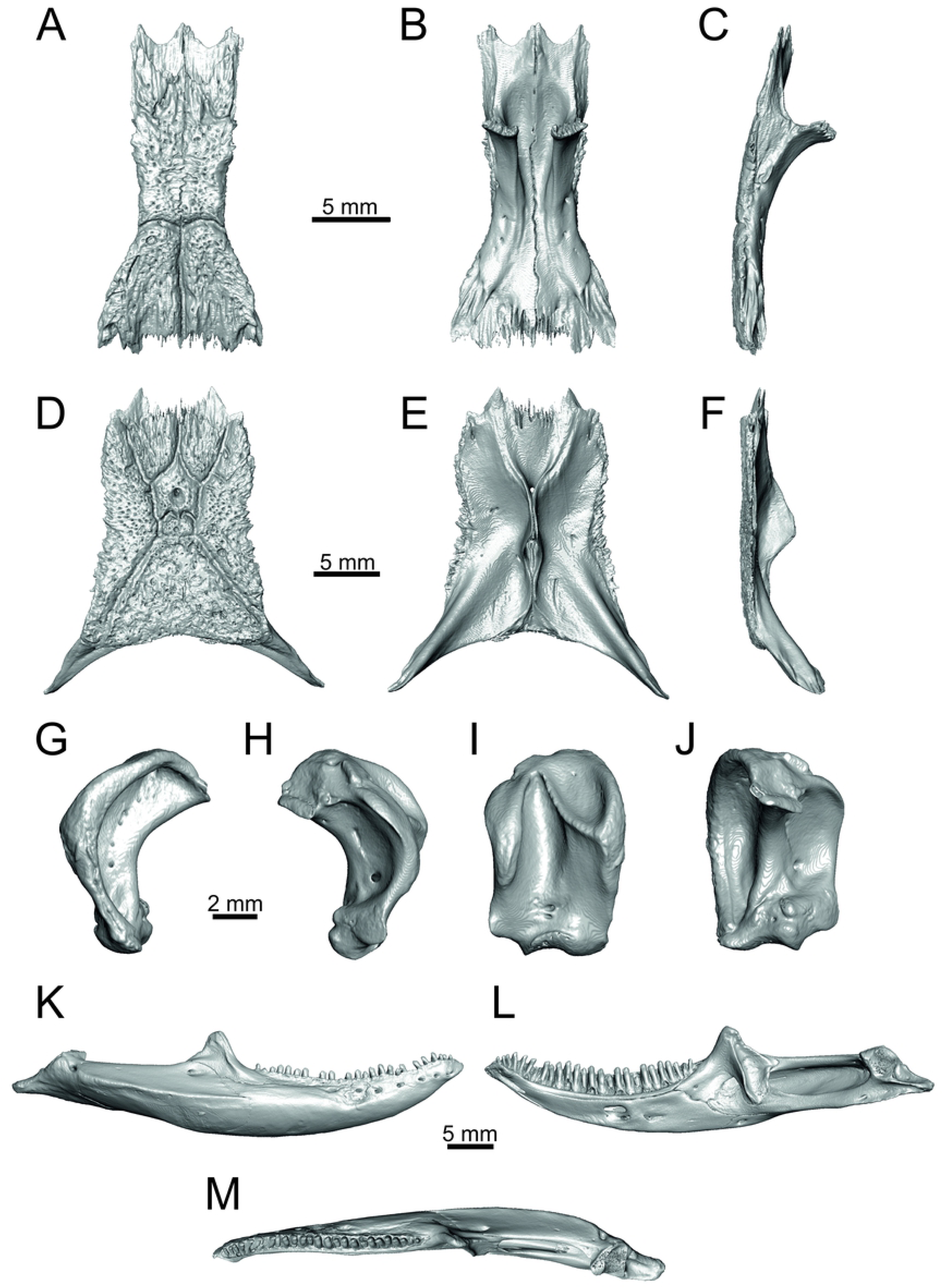
Skull and selected cranial elements of extant *Timon lepidus*. Frontal in dorsal (A), ventral (B) and lateral (C) aspects. Parietal in dorsal (D), ventral (E) and lateral (F) aspects. Quadrate in lateral (G), medial (H), anterior (I) and posterior (J) aspects. Mandibles in lateral (K), medial (L) and dorsal (M) aspects.

**Fig 29.**
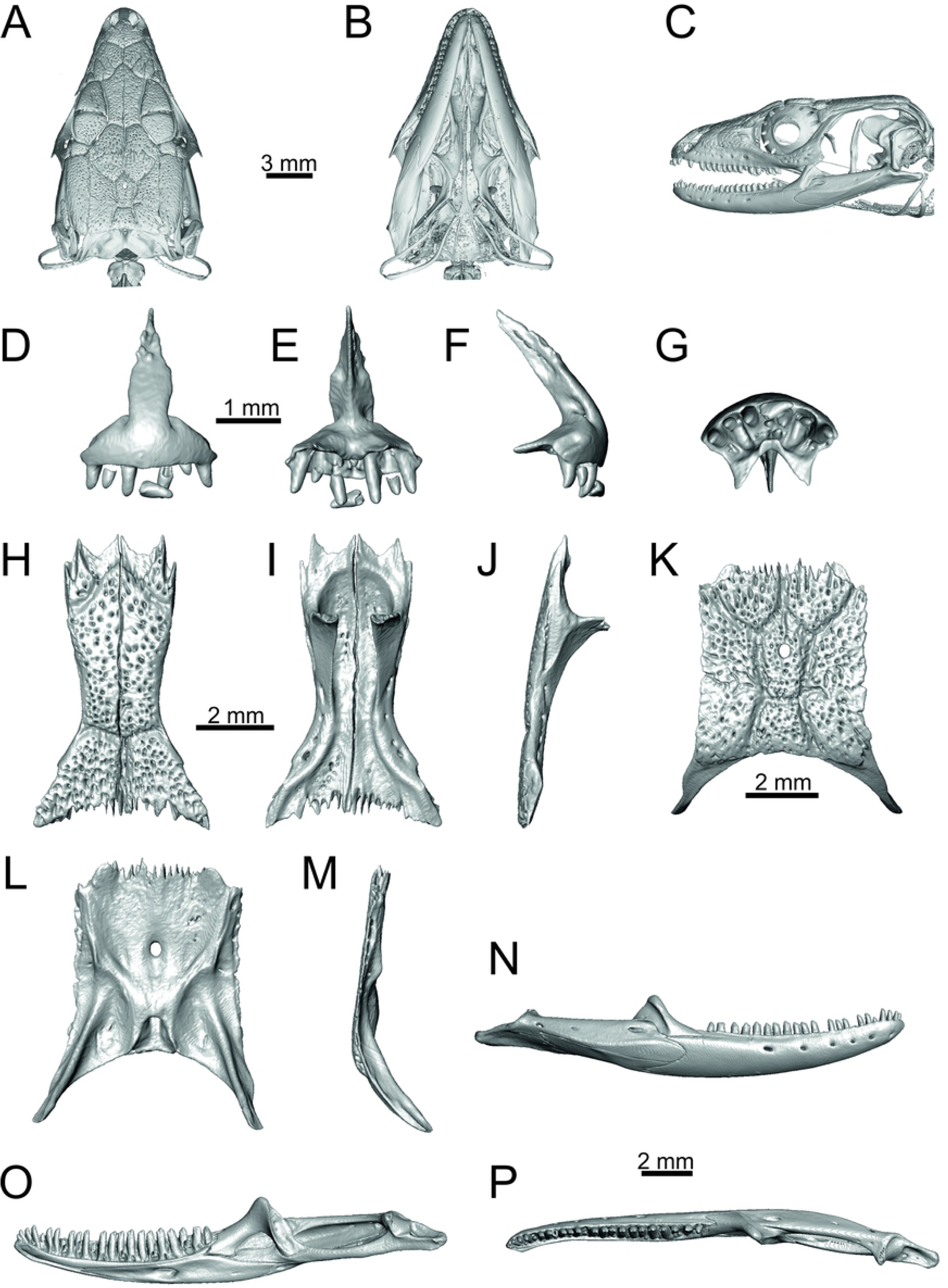
Skull and selected cranial elements of extant *Podarcis muralis*. Skull in dorsal (A), ventral (B) and lateral (C) aspects. Premaxilla in anterior (D), posterior (E), lateral (F) and ventral (G) aspects. Frontal in dorsal (H), ventral (I) and lateral (J) aspects. Parietal in dorsal (K), ventral (L) and lateral (M) aspects. Mandible in lateral (N), medial (O) and dorsal (P) aspects.

**Fig 30.**
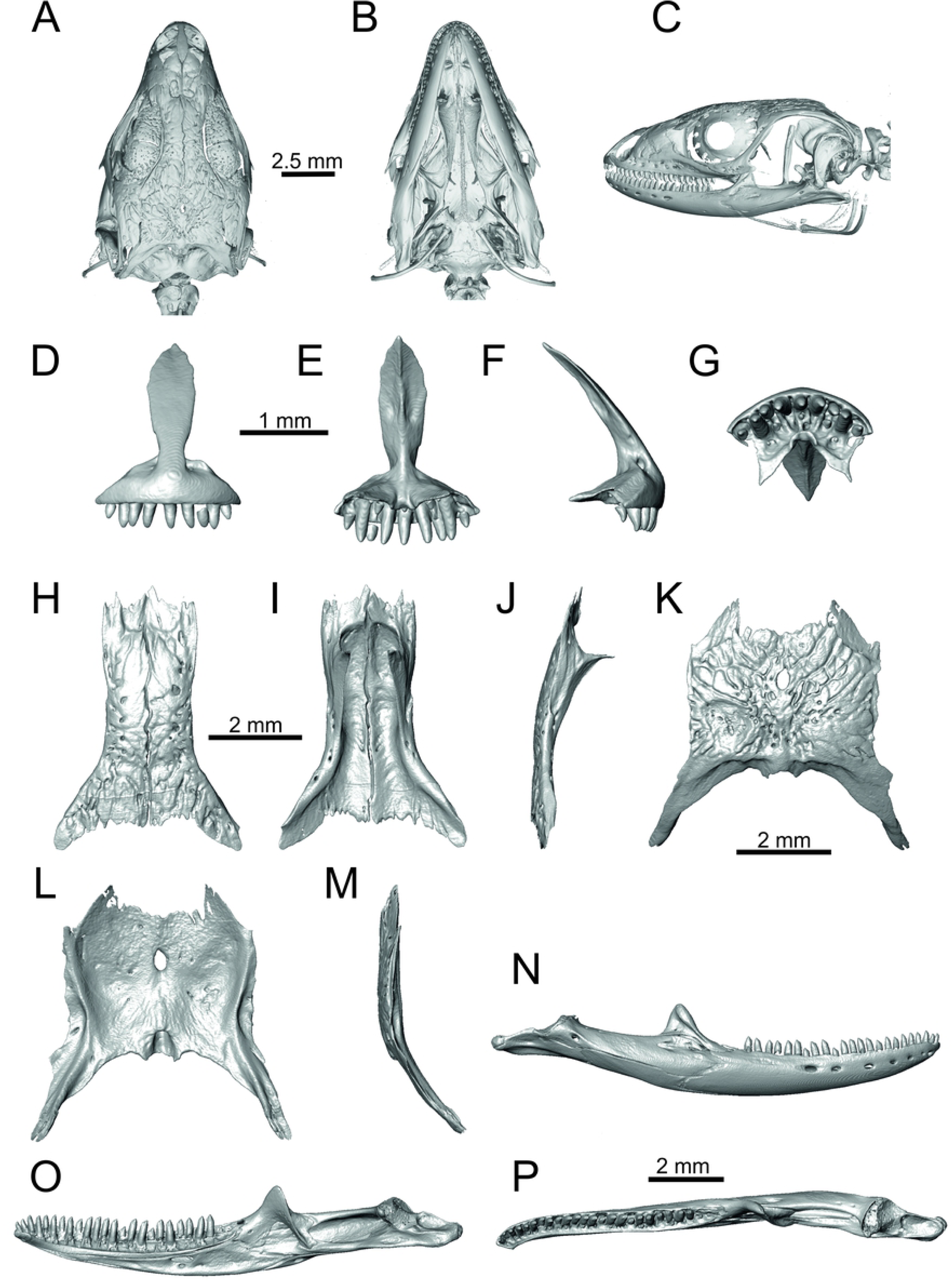
Skull and selected cranial elements of extant *Zootoca vivipara*. Skull in dorsal (A), ventral (B) and lateral (C) aspects. Premaxilla in anterior (D), posterior (E), lateral (F) and ventral (G) aspects. Frontal in dorsal (H), ventral (I) and lateral (J) aspects. Parietal in dorsal (K), ventral (L) and lateral (M) aspects. Mandible in lateral (N), medial (O) and dorsal (P) aspects.

**Fig 31.**
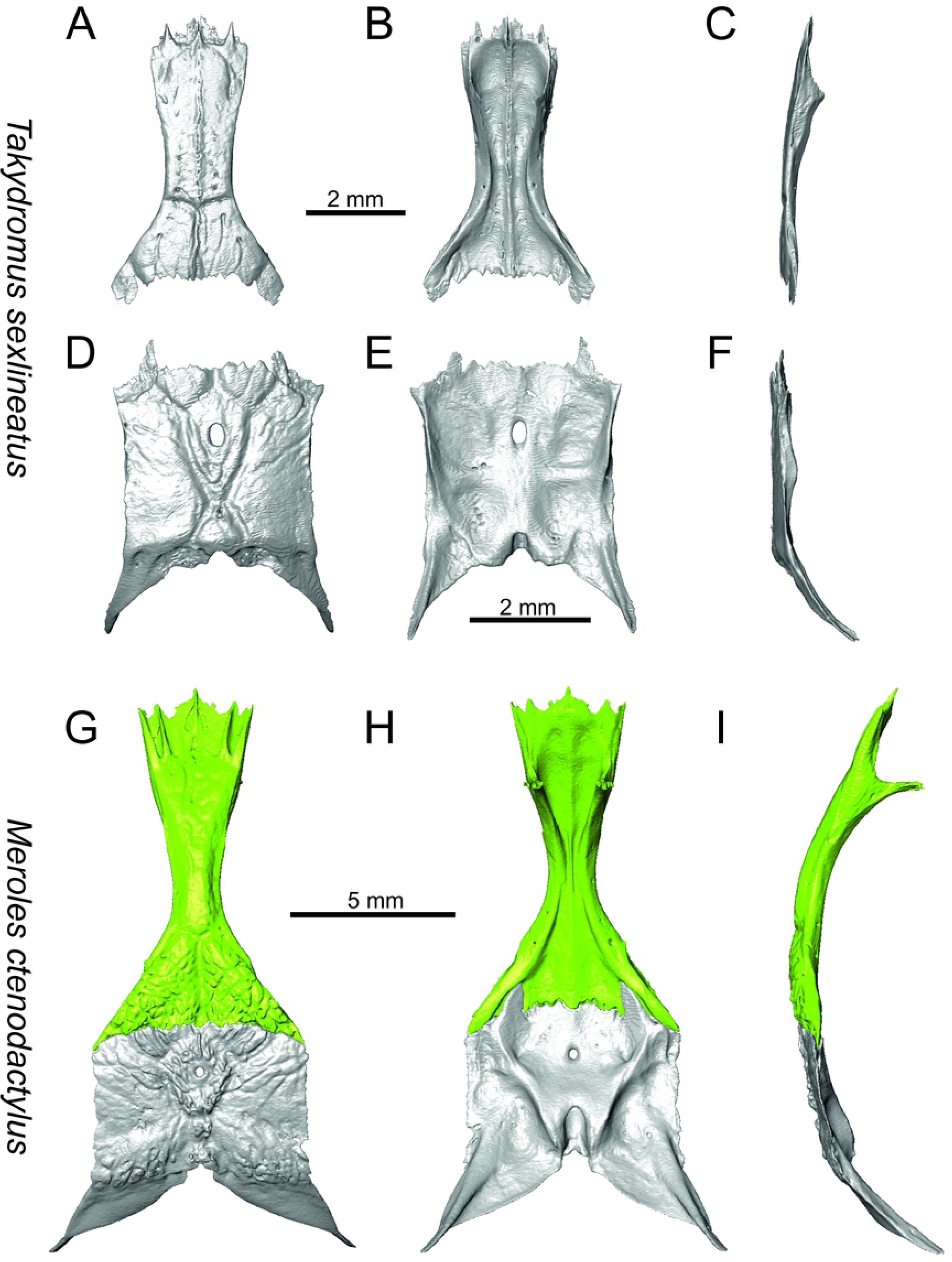
Frontals and parietals of extant *Takydromus sexlineatus* (A-F) and *Meroles ctenodactylus* (G-I) in dorsal (A, D, G), ventral (B, E, H) and lateral (C, F, I) aspects.

**Fig 32.**
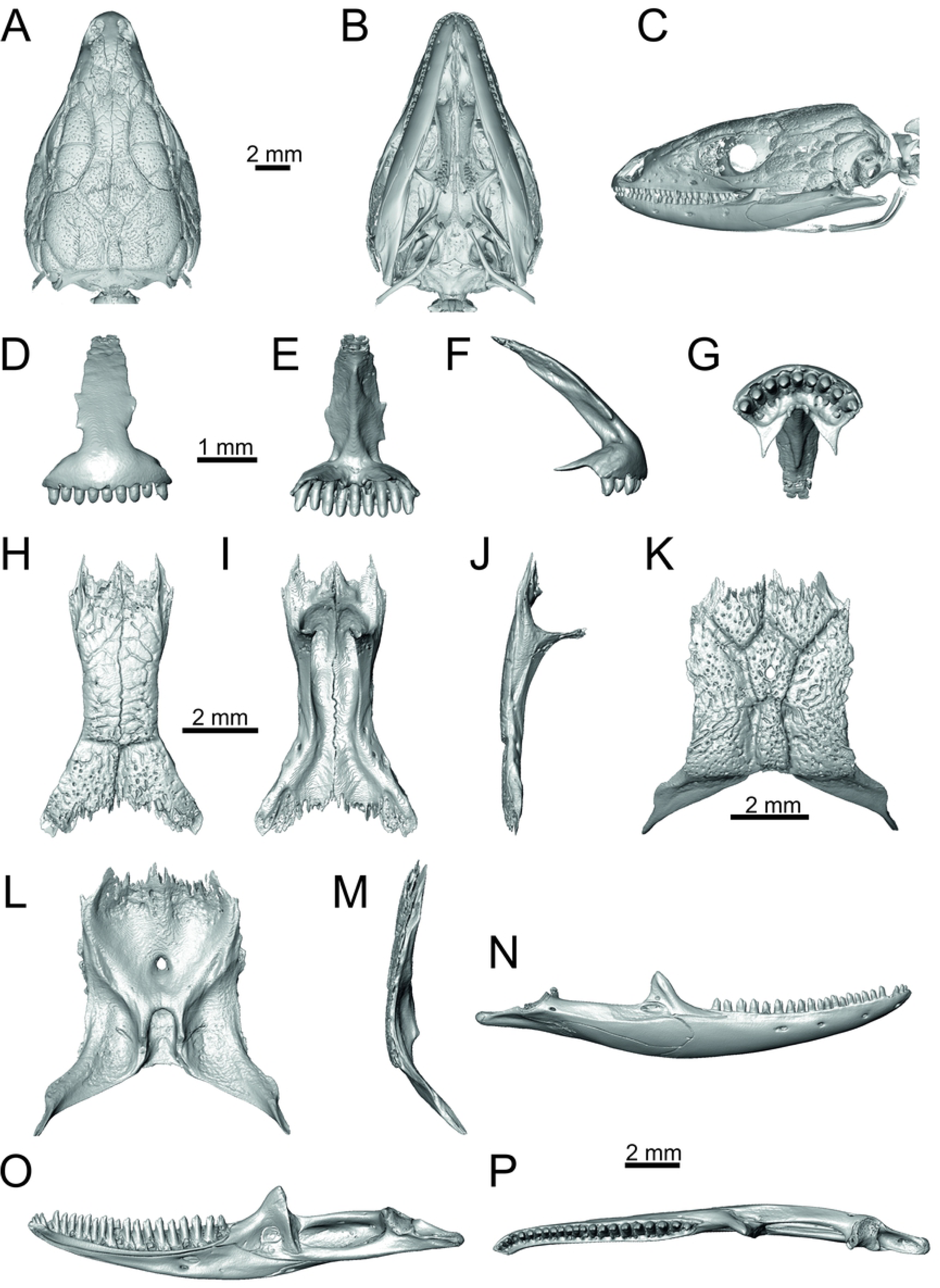
Skull and selected cranial elements of extant *Psammodromus algirus*. Skull in dorsal (A), ventral (B) and lateral aspects (C). Premaxilla in anterior (D), posterior (E), lateral (F) and ventral (G) aspects. Frontal in dorsal (H), ventral (I) and lateral (J) aspects. Parietal in dorsal (K), ventral (L) and lateral (M) aspects. Mandible in lateral (N), medial (O) and dorsal (P) aspects.

**Fig 33.**
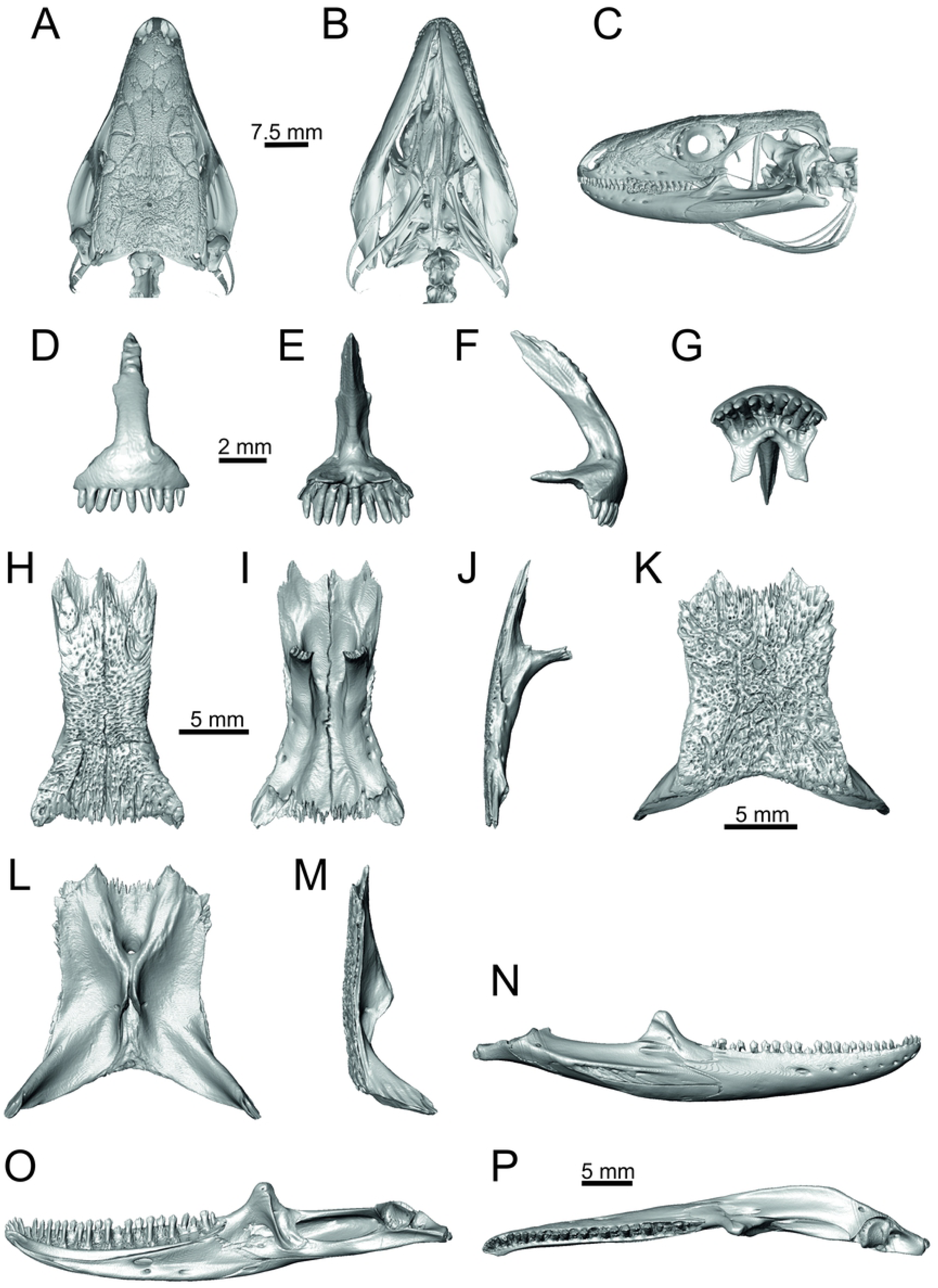
Skull and selected cranial elements of extant *Gallotia stehlini*. Skull in dorsal (A), ventral (B) and lateral (C) aspects. Premaxilla in anterior (D), posterior (E), lateral (F) and ventral (G) aspects. Frontal in dorsal (H), ventral (I) and lateral (J) aspects. Parietal in dorsal (K), ventral (L) and lateral (M) aspects. Mandible in lateral (N), medial (O) and dorsal (P) aspects.

**Fig 34.**
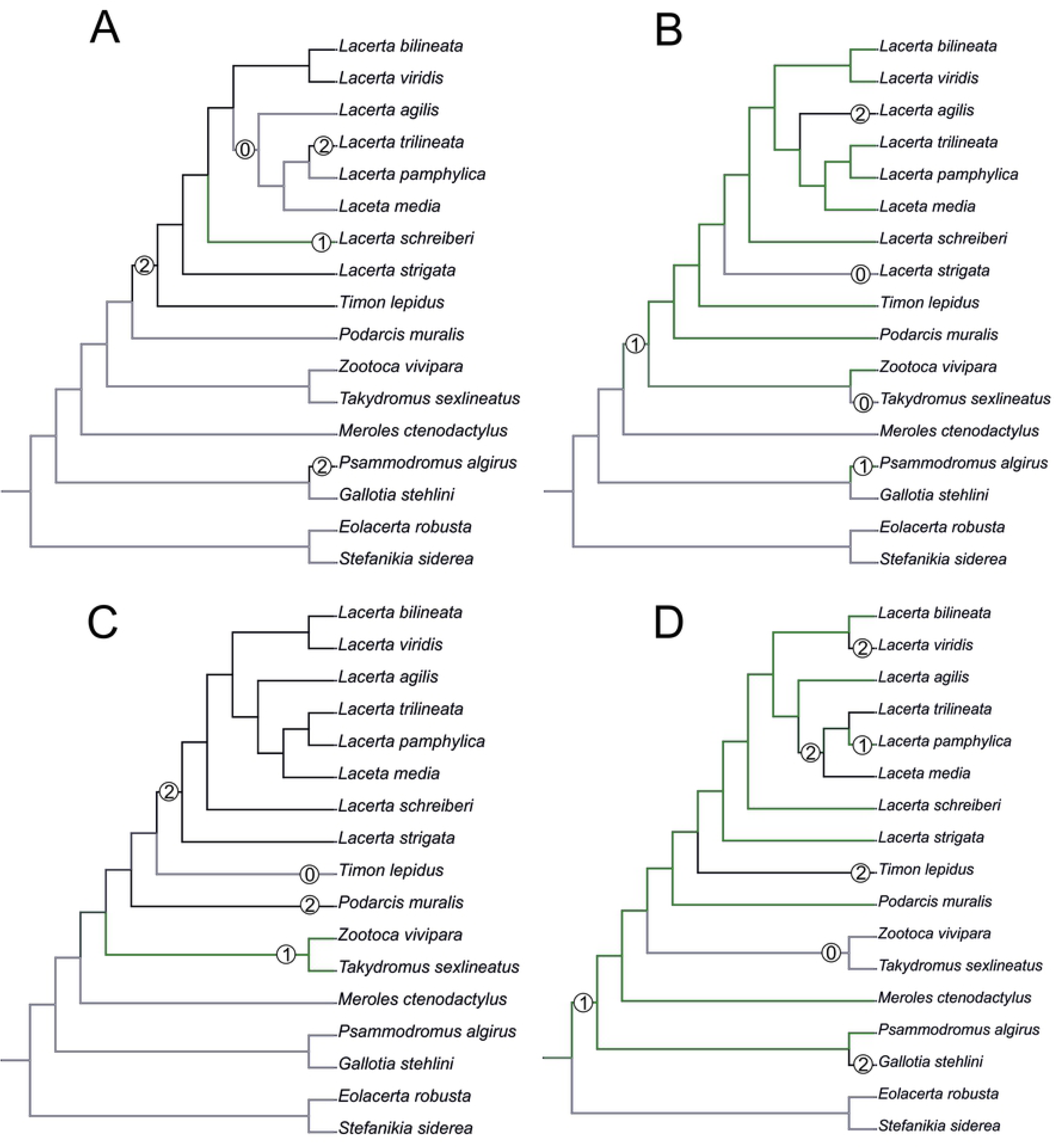
Trace character history analysis in Mesquite. (A) Cheeks covered laterally by osteoderms (absent 0, fragmented osteoderms 1, cheeks completelly covered by osteoderms 2); (B) The width of the nasal process of premaxilla (narrow 0, wide 2, extremely wide 3); (C) The stepped posteroventral process of premaxilla (absent 0,initial condition 1, present, well developed 2); (D) Parietal cranial crest medial convergence (absent, 0; tendecy to medial course present 1; crests meet medially 2.

Solnechnodolsk lacertid material can be allocated to green lizards without doubts. This can be supported by the combination of several features, e.g.:

(1) The anterior region in front of the sulcus interfacialis is long, forming more-or-less 2/3 of the anteroposterior length of the frontal. In *Timon lepidus*, this anterior region of frontal is short and the ratio of the anteroposterior length of the frontal shield vs. frontoparietal shield on frontal is approximatelly 1:1 (see [18]; or Fig. 25K here).

(2) premaxilla bears nine teeth. This is true for all green lizards and nine teeth can be observed in the Eocene taxon *Pl. lydekkeri* [8]. Among members of the clade Lacertidae seven premaxillary teeth can be found in the tribe Eremiadini, e.g. in *Acanthodactylus* or *Mesalina*, and six or seven in *Eremias persica* [47, 48]. However, such a tooth count can be observed in *Gallotia* as well [11, 41, 49], whereas in *Psammodromus* and Lacertini, the premaxillary tooth count is higher - nine or more [11]. *Gallotia* deserves a comment here. In the specimen NHMV 11031-1 (Fig. 33) the premaxilla bears nine rather than the usual seven teeth. This shows the presence of some occasional variety in this character.

(3)Parietal fossa. This structure in the posterior ventral section of the parietal forms an area where the ascending process of the supraoccipital fits into a groove (parietal fossa; see, e.g., [50]). In some taxa, the posterior margin of the parietal does not reach the anterior edge of the supraoccipital [49], leading tothe absence of a connection, which also occurs frequently in eremiadine lacertids [12]. The shape and size of the parietal fossa can be variable and can change during ontogeny (see Fig. 24 for *L. pamphylica*). In *T. lepidus*, the parietal fossa has somewhat peculiar morphology. There, the additional posterior constriction of the parietal fossa is present. This gives a lens shape to this structure (see Fig. 26B; see also [51]). Moreover, these two lineages differ in body-size [52, 53]. Similar morphology as present in *T. lepidus* can be seen in Eocene *Pl. lydekkeri*. This taxon is also similar by having fragmented parietal shield [8, 54]. Very unique morphology can be seen in the Oligocene durophagous lacertid specialist *Dracaenosaurus croizeti*, where the fossa is absent. In contrast, a parietal flange is present here [12].

(4) Medial expansion of the frontal process of the postfrontal, forming a broad area with a contact to frontal, is present. This is present in all green lizards, but is not unique to them [41]. It is also present in *T. lepidus* or *G. stehlini*, but absent in *Po. muralis, Z. vivipara, T. sexlineatus* and *M. ctenodactylus*.

### General comparison of green lizards

In the following section, osteological differences which cannot be observed in the Solnechnodolsk fossils, but are present among green lizards, are discussed:

(1) Lateral expansion of cheeks in adults. In dorsal view, *Lacerta viridis* and *L. trilineata* have distictly laterally expanded cheeks relative to other taxa (see Figs. 9, 10). This is also present in *Timon lepidus* (Fig. 27A).

(2) Osteodems covering cheeks laterally (see Fig. 9, 10). In lateral view, cheeks are not covered by osteoderms in *L. agilis, L. pamphylica* and *L. media*. The example of L. papmphylica shows that it is stable during ontogeny (see Fig. 21C, F, I). Although in adults, some small isolated osteoderms appear directly posterior to jugal in contrast to a juvenile, these do not cover the cheek. The cheeks are strongly covered by osteoderms in *L. viridis, L. bilineata, L. strigata* and *L. trilineata*. Fragmented discontinuous osteoderms in cheeks are present in *L. schreiberi*. The cheeks are strongly covered by osteoderms in *T. lepidus* (see Fig. 27C), in contrast to *Podarcis muralis* (Fig. 29C) and *Zootoca vivipara* (Fig. 30). In Gallotiinae, both states are present. Cheeks are strongly covered by osteoderms in *Psamodromus algirus* (Fig. 32C), but they are absent in *Gallotia stehlini* (Fig. 33C). The character optimization in Mesquite supports the absence of the osteodermal covering on cheeks as being the condition at the basal node of the clade Lacertidae. Character optimization in Mesquite (Fig. 34A) shows that the presence is the condition at the basal node of the clade *Timon* + *Lacerta*, whereas the covering the cheeks in *Ps. algirus* (Gallotiinae) evolved independently. Its absence in *L. agilis, L. pamphylica* and *L. media* is regarded as reversal.

(3) In *L. agilis*, the sulcus which separates the 2nd and 3rd supraocular osteoderms, meets the frontal laterally at the level of (or slightly posterior to) the sulcus interfacialis (see Fig. 9A). In all other green lizards (even in a juvenile specimen of *L. pamphylica*, see Fig. 21A) as well as in *T. lepidus, Po.muralis, Z. vivipara, Meroles ctenodactylus, Takydromus sexlineatus, Ps. algirus* and *G. stehlini*, the sulcus on the supraoculars meets the frontal anterior to the sulcus interfacialis. Thus, the sulcus interfacialis lies posterior to the region of the contact of the 2nd and 3rd supraoculars.

(4) In dorsal view, the supraoccipital in adults is highly exposed (its posterior section is not overlapped by parietal) in *L. agilis, L. bilineata, L. pamphylica, L. strigata, L. schreiberi* (see Figs 9, 10). Only a small posterior portion of the supraoccipital is visible in *L. viridis* and *L. media*. In *T. lepidus* (Fig. 27A) and *G. stehlini* (33A), the supraoccipital is completely covered by the parietal in dorsal view. In contrast to that, it is highly exposed in *Po. muralis* and *Z. vivipara*. Regarding the posterior region of the parietal, the *Z. vivipara* + *T. sexlineatus* clade can be characterized by the presence of a smooth area of the parietal table posterior to the ornamented region and by the presence of a short bilobed, posteriorly located process in the posterior mid-region.

(5) The width of the supratemporal processes (Figs. 19, 20): Broad supratemporal processes can be found in *L. media*, whereas they are rather narrow in *L. schreiberi* and *L. agilis*. In other green lizard taxa, the width spans to moderate range of width as well as in *T. lepidus* (Fig. 28D, E). Thus this state appears to be plesiomorphic among green lizards. The most ventral inclination of the supratemporateral process among lacertids studied here is present in *G. stehlini* (Fig. 33K, L).

(6) In lateral view, the skull is slightly depressed in the preorbital region, so the premaxillary region is slightly stepped (Fig. 9, 10) in *L. viridis* (marked by arrow in Fig. 9F), *L. pamphylica* (even in juvenile; Fig. 21C), *L. strigata, L. schreiberi, L. trilineata* and *L. media*. In contrast, the dorsal margin of the preorbital region is rounded in *L. agilis* and *L. bilineata*. This preorbital depression is present in *T. lepidus* (Fig. 27C). The depression is absent in *Ps. algirus* (Fig. 32C) and weakly developed in *G. stehlini* (Fig. 33C).

(7) The position of the anterior mylohyoid foramen on the splenial relative to the anterior inferior alveolar foramen. The anterior mylohyoid foramen is located ventrally to the anterior inferior alveolar foramen, but its position varies (Figs. 25, 26): it is more anteriorly located from the vertical mid-plane of the alveolar foramen in *L. strigata*, whereas it is posterior to the vertical mid-plane of the alveolar foramen in *L. bilineata, L. trilineata, L. media, L. pamphylica, L. schreiberi*. It is located directly in the vertical mid-plane of the anteroposterior length of the alveolar foramen in *L. viridis* and *L. agilis*. As for the outgroup, it is located in the posterior section in *T. lepidus, Po. muralis*, and *G. stehlini*, but in the middle in *Z. vivipara* as well as in *M. ctenodactylus*. It is located anteriorly in our specimen of *Ps. algirus*.

(8) The position of the posterior mylohyoid foramen on the angular relative to the top of the dorsal process of coronoid (Figs. 25, 26): posterior to the top of the dorsal process in *L. agilis, L. viridis, L. strigata*. It is located at the level of the dorsal process of coronoid or only slightly posterior to it in *L. bilineata, L. pamphylica, L. schreiberi, L. trilineata* and *L. media*. This second character state is also present in *T. lepidus*, whereas it is posteriorly located in *Po. muralis*and *Z. vivipara*, but anterior to the dorsal process of coronoid in *G. stehlini*.

(9) Among green lizards, the dentary of *L. media* in dorsal aspect (Fig. 26L) is markedly narrower relative to the robust postdentary mandibular region. Moreover, the articular + prearticular element forms a large mediolateral keel (or flange; it is exposed even in lateral aspect, see Fig. 26J). Although the posterior region is more robust relative to the dentary region in *T. lepidus*, this keel is not developed here (Fig. 28K-M).

### The Miocene Solnechnodolsk fossils compared to the extant taxa

Today, the area to the east of the Black Sea (where Solnechnodolsk is located) is occupied by *Lacerta agilis, L. strigata* and *L. media*; whereas *L. viridis* is distributed in northern, western and southern regions (e.g., Turkey) of the the Black Sea [21, 55, 56]. Nowadays, *L. trilineata* can be found in southern and western areas around the Black Sea [21]. The lacertid material from the upper Miocene of the Solnechnodolsk locality shares a combination of a features that is found only in *L. trilineata* among extant species. However, caution is needed here, because the sample size here does not fully allow to discuss the patterns of ontogenetic and sexual variation that must be present in these lizards and which would introduce more variation. For this reason, we decided to allocate this material as *L*. cf*. trilineata*. This combination of features is as follows:

#### Premaxilla

(1) The nasal process of premaxilla is moderately wide, gradually expanding laterally from its base in a posterodorsal direction. This is identical to that of *L. trilineata*. The process is very broad in *Lacerta agilis*, whereas it only slightly expands laterally in posterodorsal direction (although small variation can be present), being thus rather narrow (relative to others) in *L. viridis, L. strigata, L. pamphylica*, or *L. media* (see Figs. 11–12). The condition in *L. schreiberi* is similar to that of *L. trilineata*. However here, the base of the process is already slightly expanded laterally, so the lateral margins of the mid-region of the process are slightly concave (a mid-constriction is present here above the base of the nasal process, rather than gradual expansion from the base; see Fig. 12). In *Timon lepidus*, the nasal process of the premaxilla is broad in the ventral region, but is posterodorsally stepped - the posterodorsal termination is suddenly narrow (see Fig. 27D, E). A distinctly thin posterodorsal process, which gradually narrows posterodorsally, can be observed in *Gallotia stehlini* (here, a small lateral step can be present at the half-way point of the posterodorsal length of the process, see Fig. 33D-E). Moreover, the maxillary processes are reduced in this taxon [11]. The narrow nasal process of the premaxilla is present in members of the Eolacertidae [9]. The broad process among lacertids can be therefore most likely regarded as derived. Character optimization in Mesquite evaluated this change in two equally parsimonious ways (Fig. 34B): as the condition at the basal node in the the tribe Lacertini with an additional reversal in *T. sexlineatus* and, independently, *L. strigata*, or as the condition at the basal node of the clade formed by *Podarcis* + *Timon* + *Lacerta*. The wide nasal process in *Psammodromus algirus* (it is narrow in *Ps. hispanicus* [41]) represents an independent derivation.

(2) The vomerine processes (formed by the posteriorly expanded supradental shelf of the premaxilla) are moderately long. They are almost absent, not posteriorly expanded in *L. bilineata*, whereas extremely long processes are present in *L. pamphylica*.

#### Maxilla

(3) The posteroventral process of the maxilla is stepped as it is in all green lizards studied here (Fig. 13–14). However, the notch between the dorsal portion and ventral portion is only weakly developed in the Solnechnodolsk fossil. A similar condition is present in *L. trilineata* and *L. agilis*. However, the notch appears to be fully absent in *L. strigata*, and *L. schreiberi*. A strong notch is developed in *L. bilineata, L. pamphylica* and *L. media* (in these two last taxa, the dorsoventral height of the termination of the posteroventral process is large, see Figs. 13K, 14J; for *L. media*, see also [57]). This character state appears to be stable in all studied specimens of *L. trilineata* and *L. pamphylica* (juvenile and adults, see Fig. 21). However, *L. viridis* deserves a comment here. The notch is almost absent in NHMV40137 (see Fig. 13D, E), but is present in other specimens, as DE 51 (Fig. 13G; for other see see Digimorph.org [58]). Therefore it seems to be reasonabe to expect that usually the notch is well-developed in *L. viridis*. In contrast, the termination is not stepped in *T. lepidus* (Fig. 27H, I) and Eocene *Pl. lydekkeri* [8] and it is absent in *Meroles* or *Eremias* [48]. The termination is not stepped in the Paleogene clade Eolacertidae [9, 59]. Thus the presence of this step appears to be a derived character state among Lacertidae. The character optimization in Mesquite (see Fig. 34C) supports the presence of a stepped process as being the condition at the basal node of the green lizards, whereas this character state in *Po. muralis* evolved independently. The tendency to form a dorsal step apperas to be present in the basal node of the tribe Lacertini.

(4) The posteroventral process rises anteriorly, gradually continuing to the nasal process. Where the two processes contact, only a small dorsal curvature is present (it indicates the posterior end of the nasal process). In *L. media*, the posteroventral process is of almost uniform height along its length, only slightly higher anteriorly than posteriorly. The contact with the nasal process in this species is clear, the posterior margin of the ventral portion of the nasal proces is almost vertical to the posteroventral process of maxilla. In *L. pamphylica*, the posteroventral process is also uniformly deep and its dorsal margin flows anteriorly into the nasal process, the two forming a smooth concave margin.

(5) The presence of well separated osteoderms fused to the lateral side of the ventral portion of the nasal process of maxilla and the ratio of the anteroposterior length of the anterior osteoderm relative to the posteriorly located sculptured region. The anteriorly located osteoderm is large. The sulcus which separates the anterior osteoderm from the posteriorly located sculptured region, has a posteroventral course virtually pointed to the 4th labial foramen (counted from anterior). The anteroposterior length of its ventral margin forms 1/3 of the entire ventral margin of the sculptured region. Such a condition is present in *L. trilineata*, but not in *L. media, L. schreiberi, L. pamphylica, L. bilineata*. The separation of the anterior osteoderm by a sulcus is present in *L. strigata and L. viridis* as well. However, the anterior osteoderm in these species is distictly smaller in comparison to the dominant posterior osteoderm. In *L. agilis*, there is an absence of fused osteoderms (see Fig. 13A; see also [60]). In *T. lepidus*, the anterior osteoderm is small (Fig. 27J).

(6) The last labial foramen is located at the level of the 7th tooth position. This is present is adults of *L. trilineata, L. viridis* and *L. bilineata*. It is located at the level between of 4th and 5th tooth position in adults of *L. strigata*, 6th in *L. agilis*, 8th in *L. schreiberi*, between 8th and 9th in *L. pamphylica* and 10th in *L. media*. It should be noted that this character can be different in at least some juveniles, because the posteriormost labial foramen is present at the level of the 6th position in a juvenile specimen of *L. pamphylica* studied here.

(7) The septomaxillary (internal) ramus of the premaxillary process of maxilla is larger and more robustly developed than the external one. This condition is present in *L. trilineata, L. schreiberi, L. media* and *L. agilis*. The oposite condition can be found in *L. bilineata*and *L. strigata*. These two rami are almost equally developed in *L. viridis* and *L. pamphylica*. In *T. lepidus*, the septomaxillary ramus of the premaxillary process is more robust relative to the external one.

#### Jugal

(8) The angle between the central line of the posteroventral and postorbital process of jugal is around 78°. It is around 75° in *L. bilineata*. 70° can be found in *L. media* and *L. viridis*, whereas it is 67° in *L. strigata* and *L. agilis*, 65° in *L. pamphylica* and 60° in *L. schreiberi*. It is 60° in *T. lepidus*. It should be noted that the morphology of Solnechnodolsk jugal slightly resembles that in *L. bilineata*. However the posteroventral process in Solnechnodolsk specimen appears to be better defined, not having such concave dorsal margin. In *L. bilineata*, the posteroventral process appears to be shorter, less defined and bluntly ended if compared to that in other green lizards (Fig. 15E-F; see also [61]). A condition similar to that seen in the Solnechnodolsk fossil is present in *L. schreiberi* (Fig. 16C-D), but here the posteroventral process of the jugal is narrow and more dorsally inclined (slightly dorsally inclined process is also present in *L. media*). In *T. lepidus*, the posteroventral process is broad and well developed (see Fig. 27K-L).

(9) Width of postorbital process relative to posteroventral process: The postorbital process is wide. The wide postorbital process is present in *L. trilineata, L. viridis, L. bilineata*, but is somewhat narrow in *L. schreiberi, L. strigata, L media, L. pamphylica*. In *L. agilis*, the postorbital process is moderately narrow if compared to others. In *T. lepidus*, the postorbital process is wide.

#### Frontal

(10) The length of the anterolateral process relative to the anteromedial process. Although only the base of the anteromedial process is preserved, it can be estimated that it did not reach the level of the anterior end of the anterolateral process, thus was shorter in comparison to the anterolateral process. This is identical to that of *L. trilineata*. The relative length of these two anterior processes varies among species: a) the anterolateral processes are long whereas the anteromedial ones are indistinct in *L. viridis* and *L. pamphylica*; b) the anteromedial process is developed, but shorter than anterolateral one in *L. schreiberi, L. trilineata* and *L. media*; c) both reach more-or-less the same level anteriorly in *L. agilis, L. bilineata, L. strigata*. In *T. lepidus*, both processes reach approximatelly the same length. The same condition is present in *Po. muralis*. In *Ps. algirus* and *G. stehlini*, the lateral processes reach the further anteriorly than the medial ones.

(11) The lateral mid-constriction of the frontal. The position and level of the constriction is identical to that of *L. trilineata*. The most pronounced mid-constriction of the frontals is present in *L. strigata* and a distinct constriction can be also found in *L. media*. The widest frontals relative to their antero-posterior length is present in *L. schreiberi* and *L. bilineata*. In *Timon lepidus* and *Zootoca vivipara*, frontals are wide, however, they are constricted in *Podarcis muralis*. The marked constriction can be seen in *Meroles ctenodactylus* and a similarly strong constriction appears to be common in members of the tribe Eremiadini (for *Acanthodactylus erythurus*, see [41]).

#### Parietal

(12) In the dorsal surface of the parietal, the interparietal shield is pierced by the parietal foramen in the anterior half of its anteroposterior length. This is identical to *L. trilineata*. In all other green lizards (for all green lizards, see Figs. 19–20), the position of the parietal foramen is located in the posterior half of the interparietal shield (*L. agilis, L. strigata, L. schreiberi, L. media, L. pamphylica*), or more or less in the mid-region (*L. viridis, L. bilineata*). In *L. pamphylica, L. schreiberi, L. trilineata* and *L. viridis*, the interparietal shield is entirely restricted to the parietal by the frontoparietal shields that meet to form a medial suture that interpolates between the anterior margin of the parietal shield and the suture with the frontal bone. In *L. agilis* and *L. bilineata* the frontoparietals are in point contact, while in *L. media*, the frontoparietal shields do not meet on the parietal bone, allowing the anterior margin of the parietal shield to contact the suture with the frontal, and in *L. strigata*, a small anterior portion of interparietal shield continues on the posterior mid-region of frontals. In *T. lepidus*, the interparietal shield is pierced in its anterior region (Fig. 26A; see also [51]). An even more anterior location of the foramen can be observed in *Po. muralis, Takydromus sexlineatus*, and *G. stehlini* (as well as *Janosikia ulmensis* [11]). The interparietal shield is pierced more-or-less in the middle in *Z. vivipara* and the forman is located in posterior region of the interparietal shield in *Ps. algirus*.

(13) The occipital shield of the parietal in *L. trilineata* is large, occuping the largest area of the parietal table among all green lizards (although it should be noted that the anteroposterior length of the occipital shield is smaller than the length of the interparietal shield). The occipital shield is broad, wider than the interparietal shield. The lateral margins of the occipital shield are convex laterally rather than having a straight course. The same condition is present in the Solnechnodolsk lacertid. Among the taxa studied here, the occipital shield is large in *T. lepidus* (but also in *Pl. lydekkeri* [8]) and *G. stehlini* (for this character state see also [11]).

(14) The lateral margins of the parietal are not completelly preserved and can be only estimated. However, they appear to have a lateral concave course similar to *L. media, L. pamphylica, L. trilineata* and *L. viridis*. In *L. trilineata* and *L. media*, the small mid-constriction is present and the posterior portion is equally wide as the anterior portion. Such constriction is also present in *L. viridis* and *L. pamphylica*, but here the posterior portion is slightly wider than the anterior one. In *L. agilis* (slightly concave margins) and *L. bilineata* (slightly convex margins), the whole parietal narrows anteriorly. In *L. schreiberi*, the lateral margins are more-or-less straight. In *L. strigata*, the lateral margins run lateraly from each other anteriorly, thus the anterior portion is slighty wider than the posterior one. In *T. lepidus*, the parietal table is anteroposteriorly long and slightly narrows anteriorly. The lateral margins of the parietal table are slightly concave in this taxon.

(15) On the ventral surface of the parietal, the parietal cranial crests converge strongly posteriorly, forming a distinct median crest slightly posterior to the parietal foramen (for this character see [23]). This is identical to adults of *L. trilineata* and *L. viridis* and slightly similar condition can be observed in *L. media*. The diffence here between *L. trilineata* and *L. viridis* is in the course of the parietal crest. In *L. viridis*, the crest on each side continues anterolaterally in a more or less straight course. Thus, the virtual line extending anteriorly from the posterior section of the branch would meet the anterolateral corner of the parietal (see also [9]). In *L. trilineata*, the parietal crest on each side is angled approximately at the level of the anterior margin of the parietal foramen. Here, the parietal crest turns more anteromedial relative to its posterior section. Thus, the virtual line extending anteriorly from the posterior section (posteromedial to this angle) would not meet the corner of the parietal table, but would finish on its anterolateral side. This is identical to the condition present in the Solnechnodolsk parietal GIN 1145/283 (see Fig. 5B). In other green lizards, an evident tendency to a medial course of the parietal crest is present, but not in the distinct form as it is in *L. trilineata* and *L. viridis*. Moreover, this character state changes during ontogeny. It is absent in juveniles of green lizards, here supported by *L. pamphylica* (see Fig. 21J). The parietal crests converge in *T. lepidus* and *G. stehlini*. Only a tendency to more-or-less medial course can be observedin adults of *Po. muralis, M. ctenodactylus* and *Ps. algirus*. The tendencey to a more or less medial course is absent in *Z. vivipara* and *T. sexlineatus*, and in known Paleogene members of the clade Eolacertidae (it is regarded here as absent, but members of this clades might appear to show somewhat an initial tendency [9]). The juveniles of green lizards therefore show the morphology of their ancestral lineage and the converging of the parietal crests can be regarded as derived among the Lacertidae. The character optimization in Mesquite (Fig. 34D) supports the presence of a tendency to a medial course of the parietal crests as being the condition at the basal node of the clade Lacertidae (this is also supported by the presence of medially merged parietal crests in the Eocene *Pl. lydekkeri* [8]. Its absence in *T. sexlineatus* and *Z. vivipara* is regarded as reversal. Character optimization in Mesquite shows that the presence of the medially merged parietal crests in *G. stehlini, T. lepidus* and several members of *Lacerta* evolved independently. In the clade *L. trilineata* + *L. pamphylica* + *L. media*, character optimization in Mesquite evaluated this change in two equally parsimonious ways: as the condition at the basal node in the this lineage with an additional reversal in *L. pamphylica*, or as representing independent derivations of the medially merged crests in *L. trilineata* and *L. media*.

(16) The ventrolateral ridge in the internal region gradually disappears anteromedially. This character state can be observed in *L. trilineata, L. pamphylica, L. strigata* and *L. media*, but not in *L. agilis, L. viridis* (see also [9]) and *L. schreiberi*. Here, this ridge continues markedly almost to the parietal crests.

(17) Parietal fossa - in regard to the Solnechnodolsk parietals, we can exlude *L. media* here. This taxon is characterized by very narrow wedge shaped fossa, with an additional process located in the anterior end of the fossa (posteroventral process in Fig. 20L). This process is directed posteroventrally.

#### Postfrontal

(18) In Solnechnodolsk fossils, the postfrontal and postorbital are separated individual elements. This is present in all specimens of green lizards (e.g., *L. viridis*; see Fig. 22A) examined here except for two species. In *L. agilis*, partial fusion of these two elements has occured, with traces of the suture being easily recognized. However in *L. schreiberi*, the fusion is present in a stronger degree, where traces of the suture can be recognized only in the posterior region of the postorbitofrontal element (Fig. 22B). On its dorsal surfaces, two osteoderms are separated by a longitudinal sulcus. The fusion of the postorbital and postfrontal bones to form a postorbitofrontal can be seen in *T. lepidus* (Fig. 22C), but not in *Po. muralis* (Fig. 22D). The fusion is present in *Z. vivipara* (Fig. 22E) and in *T. sexlineatus* (Fig. 22F), but also in *M. ctenodactylus* (Fig. 22G) and in members of Gallotiinae (Fig. 22H; see also [11, 62]), as well as in the Eocene *Pl. lydekkeri* [8]. The postfrontal and postorbital are unfused in *Eremias* and *Mesalina* [48].

#### Quadrate

(19) The tympanic crest, which forms the anterior margin of the quadrate is angled approximatelly in the mid-region. This is present in *L. trilineata, L. media, L. strigata, L. viridis, L. pamphylica* and *L. schreiberi* (Figs. 23–24). This anterior margin is completely rounded in *L. agilis* and *L. bilineata*. The angle is also present in *T. lepidus*.

(20) In lateral aspects, the ventral half of quadrate (ventral to angulation of anterior margin) gradually markedly narrows ventrally. Thus this portion is much narrower than the dorsal portion. This condition is identical to that of *L. trilineata, L. viridis* and *L. media*. However in the latter taxon, the quadrate is more robustly built if compared to other taxa. This difference between the width of the dorsal and vertral portion is not so pronounced in other taxa. In *T. lepidus*, the whole quadrate is robust.

#### Pterygoid

(21) Pterygoid dentition. Teeth on the pterygoid, which are often found in species of *Lacerta, Timon, Anatololacerta, Gallotia*,and *Psammodromus algirus* [49, 62], shows several patterns of their distribution. The teeth on the Solnechnodolsk pterygoid are arranged strictly in a single line. This is present in *L. trilineata, L. agilis, L. viridis, L. strigata* and *L. media*. In *L. bilineata, L. pamphylica* and *L. schreiberi*, teeth are arranged in the elliptical area, where the central region is wider-here at least two teeth can be found next to each other (in medio-lateral plane). This second condition is also present in *T. lepidus* and *G. stehlini*. In *Ps. algirus*, teeth on pterygoid are arranged in a wide area and not in a single line. Because the clade Gallotiinae are sister to Lacertidae [6, 63], we regard this second condition as plesiomorphic. Among here studied taxa, pterygoid dentition is absent in *Po. muralis* or *Z. vivipara*.

#### Dentary

(22)Number of unicuspid teeth. In the Solnechnodolsk material, bicuspidity sometimes starts in 7th tooth, but this can be variable even in extant taxa. Kosma [64] stated that first five anterior dentary teeth are unicuspid, whereas more posterior teeth are bicuspid in both *L. viridis* and *L. trilineata*. In contrast, unicuspidity is restricted only to the first two anterior teeth in *L. agilis* [64].

(23)The position of alveolar foramen in dentary at the level of the 7th tooth position (counted from posterior). Such condition is present in *L. trilineata* and *L. schreiberi*, whereas it is present at the level of the 6th tooth position in *L. bilineata, L. strigata, L. agilis, L. viridis*. The alveolar foramen is located further anteriorly, at the level of the 8th tooth position, in *L. pamphylica* and at the level between 9th-10th tooth positions in *L. media*. Although the location of the alveolar foramen can be informative, it should be noted that the positions of the alveolar foramen in lacertids, like virtually all lizards, should not be interpreted as absolute due to its variations.

### *Lacerta* cf. *trilineata* in the late Miocene of Russia

The whole detail comparison (see above) strongly supports an allocation of the Solnechnodolsk lizard fossil to *Lacerta* cf. *trilineata*. According to Ahmadzadeh et al. [28], *L. trilineata* seems to evolved and diverged in western Anatolia during the Pliocene or early Pleistocene. Sagonas et al. [16] suggested a longer history of this taxon, with the first split occurred at the early Tortonian (9.55 Mya). Based on these authors, the split of *L. viridis* and *L.bilineata* is also dated in the late Miocene (6.78 Mya). This is in a strong contradiction with the recently published results of Kornilios et al. [27]. These authors suggested that the radiations of all major greenlizard groups, including *trilineata+pamphylica*, occurred in parallel in the late Pliocene. All three studies used the same calibration points and the same data (mtDNA) for their time estimates, but differed in their inference methods. The Solnechnodolsk material shows support of the hypothesis of Sagonas et al. [16] and forms the first potential evidence of the occurrence of the lineage of *L. trilineata* already in the late Miocene.

Green lizards form a dominant component of the squamate paleofauna from the Solnechnodolsk locality (see also [3, 33]), showing their successful adaptation to the paleoenvironment of this area east of the Black Sea during the late Miocene. In this geological sub-epoch, the climate in Europe remained warm in the MN 13, however the mean annual temperature dropped when compared to the Miocene Climatic Optimum [1, 65]; the amplitude of the temperature drop variing geographically [66, 67]. This led to a more distinct climatic zonation on the European continent and to the survival of thermophilic ectothermic animals only in southern regions [1, 3, 58, 68]. The rapid climatic changes during the Miocene (and the Cenozoic in general) most likely led to the broad radiation of lacertid lizards in Europe. In the late Miocene, major vegetation changes are documented in the southeastern areas of this continent. Based on paleobotanical data, Ivanov et al. [69] reported slight cooling and some drying at the beginning of the late Miocene, followed by cycling changes of humid/dryer and warmer/cooler conditions. The occurrence of *Varanus* sp. in Solnechnodolsk suggests a mean annual temperature not less than around 15 °C [3]. Today, *Lacerta trilineata* occupies Mediterranean-type shrubby vegetation, sandy shores, arable land, pastureland, plantations, and rural gardens. The diet of this taxon consisted mainly of insects [70, 71]. In conclusion, the material of *Lacerta* together with previously-described lizard fossils [3, 33], exhibit an interesting combination of survivors and the dawn of modern species in the Solnechnodolsk locality. Thus, this locality forms an important evidence of a transition of an archaic Miocene world to the modern diversity of lizards in Europe.

### Selected fossils in European deposits vs. extant taxa

Lacertids are often reported from the fossil record of Europe [18, 54, 72–75] and from areas adjacent to Solnechnodolsk (the Black Sea area), including lacertid material previously described from the early Pliocene of Turkey. However due to the fragmentary nature of the elements, they were allocated only to cf. *Lacerta* sp. [76]. A locality identical in age (MN 13) to the Solnechnodolsk locality is represented by the Ano-Metochi locality of northern Greece, where the clade Lacertidae has also been identified. However, due to the fragmentary nature of the material, only an allocation at the family level is supported without doubt [2]. It should be noted that a herpetological assemblage from the Romanian margins of the Black Sea, is described from slightly older sediments - MN 7+8 [77]. The lacertid material there consists only of jaws and frontal fragments and was identified as *Lacerta* sp.

In this section, previously described lacertid fossils from the European Cenozoic, which can may have strong affinities with modern lineages, are discussed. The Eocene taxon *Plesiolacerta lydekkeri* is regarded as in or close to the crown Lacertidae [8]. As was already recognized by Hoffstetter [78], this taxon reseambles morphologically the extant *Timon lepidus* [79]. These shared features (some of them are potentional synapomorphies) areas follows: (1) the shape of the nasal process of premaxilla (see above); (2) shape of the maxillary processes of the premaxilla; (3) presence of nine teeth on premaxilla; (4) posteroventral process of maxilla is not stepped; (5) the overall shape of the nasal process of maxilla; (6) the maxillary crest (the *carina maxillaris*) starts from the supradental shelf a the level of the 4th tooth position; (7) the course of the maxillary crest - its anterior section runs posteriorly almost in a horizontal level (this gives the appereance to this section as being depressed). It starts to rise posterodorsally at the level of the 6th tooth position; (8) similar ratio of the anteroposterior length of the frontal and frontoparietal shields; (9) fragmentation of the interparietal shield (= presence of the transitional shield, see [8]; although it should be noted that not all individuals of *Timon* possess this character, see e.g. [37]); (10) large occipital shield; (11) a similar type of sculpturure of osteodermal shields fused to cranial bones; (12) posterior constriction of the parietal fossa of the parietal; (13) the anteroposterior length of the parietal table greater than its width; (14) postfrontal and postorbital fused to form a postorbitofrontal; (15) markedly rounded subdental shelf of dentary; (16) the alveolar foramen on dentary located at the level of the 16th tooth position (counted from anterior) in our specimen of *T. lepidus*, whereas it is located at the level of the 17th tooth position in *P. lydekkeri*; (17) the size of the elements, indicating animals with a similar large body size.

These are not just shallow similarities, because some features e.g., additional posterior constriction of the parietal fossa, are present only in these two taxa and cannot be only a matter of a large size (this feature is not present in large members of *Gallotia*, e.g. *G. stehlini*). It seems to be more plausible that such similarities might reflect possible relationship rather than represent just homoplasies. *Plesiolacerta* is also known from late Oligocene deposits of Germany, where the species *P. eratosthenesi* is described [8]. The molecular clock estimation of the evolutionary divergence between *Lacerta* and *Timon* dates back to 18.6 Mya [80], or 20.7 [27]. However, if *Plesiolacerta* has a close relationship to *Timon*, it would show a much longer evolutionary history of lineage. In comparison with other lizard lineages, this would be not so unusual-one example is the presence of the modern genus *Ophisaurus* (or morphologically identical elements) in the late Eocene [81]. This problem could be resolved if fossils proving the existance of *Timon* lineage were recognized in deposits close to the Oligocene/Miocene boundary.

The parietal (UMJGP 204.750) from the late middle Miocene of Gratkorn in Austria, allocated to Lacertidae *incertae sedis* by Böhme & Vasilyan [75] deserves a comment here. Based on the comparative study presented here, this parietal is almost identical to *Podarcis muralis* (for *Podarcis*, see also [50]). If the allocation of this parietal to *Podarcis* were correct, it shows the presence of this taxon already in the middle Miocene. Therefore, this parietal requires a detailed revision.

## Acknowledgments

We thank the fellow members of the expeditions to the Solnechnodolsk locality from Institute of Arid zones (SSC RAS, Rostov-on-Don) and Geological Institute of the Russian Academy of Sciences (Moscow). For CT scanning of the extant taxa, we thank J. Šurka (Slovak Academy of Sciences). For English correction and advice, we are indebted to M. Hutchinson (South AustralianMuseum). This project was supported by 1/0209/18 from the VEGA Scientific Agency (A. Č.). The study was also conducted by government theme AAAA-A19-119020590095-9 (Russia) and supported by the Russian Scientific Fund 𝒩<u>o</u> 18-74-10081 (laboratory work) (to E. S.)

## Authors’ contributions

Conceptualized the study: AČ. Collected the specimens and data for geology of the Solnechnodolsk locality: ES. Analyzed the data: AČ. CT data and segmentation: AČ. All authors provided critical revision of the manuscript and approval of the article.

